# Beyond performance, competence, and recognition: Forging a science researcher identity in the context of research training

**DOI:** 10.1101/2023.03.22.533783

**Authors:** Mariel A. Pfeifer, C.J. Zajic, Jared M. Isaacs, Olivia A. Erickson, Erin L. Dolan

**Author notes:** Corresponding Author: Mariel A. Pfeifer, Department of Biochemistry & Molecular Biology, B210 Davison Life Sciences Bldg., University of Georgia, Athens, GA 30602.

## Abstract

**Background**

Studying science identity development has been useful for understanding students’ continuation in science-related education and career paths. Yet, how science contexts shape students’ science identity development, especially as students engage in research at the undergraduate and graduate level, is still largely unexplored. Here we integrate science identity and professional identity theories to investigate how research training shapes science identity. We focus on a specialized form of science identity we call science researcher identity. We characterize how the features of an individual’s research experience, and their personal characteristics interact to influence whether and how they see themselves as a science researcher. We accomplished this in two phases of qualitative research. First, we surveyed 548 undergraduate researchers about how their research experience influenced their identity as a scientist. Then we interviewed 30 early career researchers, including undergraduate, postbaccalaureate, and doctoral students, about their views of themselves as researchers and how elements of their research training shaped their views.

**Results:** Early career researchers (ECRs) viewed themselves as either science students or science researchers. How ECRs recognized themselves depended on how they viewed the purpose of their daily work and the extent to which they perceived autonomy and intellectual responsibility in their research. Individual-level factors, including research and researcher conceptions, research skill perceptions, and career intentions, influenced whether ECRs identified as science students or science researchers. ECRs also recognized themselves more or less as researchers based on contextual factors like the nature of their work, social interactions, and their perceptions of the norms within their research group and institution. ECRs considered how individual and contextual factors affected their science researcher identity through a lens we call ‘sense-making.’ We further detail the processes ECRs use to make sense of their science identity and the factors that influence it.

**Conclusions:** We synthesized our findings to form a conceptual model of science researcher identity development, which hypothesizes relationships among constructs related to science identity and professional identity development. Our results advance theory related to science identity, offer avenues for future investigation, and inform efforts to promote science researcher identity development.

## Introduction

Our identities reflect how we see ourselves and shape how we experience, interpret, and make sense of life events (Oyserman, 2001). The formation and development of a science identity – or the process of seeing oneself as a scientist – has been a focus of extensive research, especially research that aims to understand the experiences of students from backgrounds that are marginalized and minoritized in science. Groundbreaking research from Carlone and Johnson (2007) developed a model of science identity through an investigation of the undergraduate and graduate science education experiences of 15 women of color. This research revealed that individuals who endorsed a scientist identity highlighted the importance of **performing science** – meaning they used the tools and language of science, **feeling competent** – meaning they perceived themselves as knowing and understanding science content, and **being recognized** – meaning they saw themselves and perceived that others saw them as a science person (Carlone & Johnson, 2007). This performance-competence-recognition model has since informed the study of scientific identity development, including how particular science learning experiences shape the extent to which students identify as scientists or not.

Research experiences at the undergraduate and graduate level are important contexts for attracting and preparing the next generation of scientists (Gentile et al., 2017), including fostering students’ development of a science identity (Chemers et al., 2011; Estrada et al., 2011; Robnett et al., 2015). Undergraduates who participate in research report learning to think and work like a scientist (performance), increased confidence in their ability to carry out research tasks (competence), and stronger identification as a scientist after participating in research (Adedokun et al., 2013; Estrada et al., 2011; Frantz et al., 2017; Hernandez, Hopkins, et al., 2018; Robnett et al., 2015; Thiry et al., 2011). Furthermore, undergraduate researchers who report a robust science identity are more likely to continue into graduate education and careers involving science research than individuals who question their science identity (Estrada et al., 2018). Yet, little research has examined how students’ science identity evolves as undergraduates go onto more intensive research training that is integral to the preparation of early career researchers (Gentile et al., 2017).

In addition, a growing body of research has revealed that research experiences can vary in ways that have the potential to disrupt students’ science identity (Camacho et al., 2021; Goodwin et al., 2022; Limeri et al., 2019; Remich et al., 2016; Rodriguez et al., 2022; Thiry et al., 2012; Tuma et al., 2021). For instance, students vary in the extent to which they are epistemically involved in, or intellectually responsible for, their research (Burgin et al., 2012). Those with limited epistemic involvement may perceive themselves as having fewer opportunities for performance, having less competence, and being underrecognized as a science person. Interactions with research mentors can also influence early career researchers’ science identity development (Alston et al., 2017; Atkins et al., 2020; Gentile et al., 2017; Robnett et al., 2018). Moreover, undergraduate researchers who are mentored by both a faculty member and a graduate student or postdoctoral researcher identify more strongly as a scientist than those with only one mentor (Aikens et al., 2016).

Hazari and colleagues (2010) further developed Carlone and Johnson’s (2007) model through an investigation of undergraduates who had taken at least one high school physics course. Hazari and colleagues (2010) determined that physics interest – or the “desire or curiosity to think about and understand physics” (Hazari et al., 2010, p. 982) – and recognition by others as a “physics person” also predicted students’ identifying as a physics person, resulting in a refined science identity model of **interest**, **performance-competence**, and **recognition by others** (reviewed in Potvin & Hazari, 2013). The refined model has been used to study identity in other STEM disciplines, including engineering, mathematics, marine science, and computing (Ambrosino & Rivera, 2022; Cribbs et al., 2015; Dou et al., 2019; Godwin et al., 2016; Mahadeo et al., 2020). In studies of math and physics identity, students’ performance-competence beliefs along with interest and recognition by others are predictive of their science disciplinary identities (Cribbs et al., 2015; Godwin et al., 2016). Performance-competence is considered an integral component of science identity because students must display performance-competence to achieve recognition by others and to consider themselves to be interested in science and science-related subjects (Cribbs et al., 2015; Godwin et al., 2016). These and other studies of interest development indicate that science experiences within and outside science classrooms are critical for interest development (Alexander et al., 2012; Harackiewicz et al., 2016; Hazari et al., 2010; Hidi & Renninger, 2006). Yet, science identity models have focused primarily on understanding the self and self-views, rather than attending to how the environment, or context, influences science identity development.

Leading identity scholars have called for research on contextual factors that influence an individual’s science identity, as well as research to understand how these factors shape the evolution of one’s science identity over time (Hazari et al., 2020; Kim & Sinatra, 2018). While multiple studies show that research experiences foster undergraduates’ science identity development, at least some of this research indicates that undergraduate students already strongly identify as scientists even before they engage in research (Adedokun et al., 2013; Estrada et al., 2011, 2018; Frantz et al., 2017; Robnett et al., 2018). This raises a question of whether the current, prevailing science identity model is useful for explaining how research training experiences influence the science identity of students who already identify, to some degree, as a scientist.

Undergraduates who have engaged in research and who intend to continue research training at the postbaccalaureate and doctoral level are likely to differ in their science identity from undergraduates who have never experienced research before. Rather than identifying as a scientist or not, their professional identity is likely to be more nuanced in a way that relates to their specific education and career pursuits. Understanding this nuance is critical for informing the design and implementation of research training programs that encourage, or at least avoid dissuading, science-interested students from pursuing further education and careers in research. For instance, faculty who mentor undergraduates may misinterpret or underrecognize the science identity of undergraduate researchers, inadvertently discouraging undergraduates from continuing in research (Thompson & Jensen-Ryan, 2018). At the graduate level, such signals may be even more influential because research is central to science graduate students’ experiences. For example, doctoral students who reported that their faculty mentors prevented them from making a research presentation at a conference took this as an indication that their work was unworthy of attention (Tuma et al., 2021). Research is needed to elucidate identity development among early career researchers (ECRs) given the importance of research training for preparing the next generation of science researchers, the critical role of identity development in shaping students’ career decisions, and results indicating that research training varies in ways that could hinder students’ professional identities.

Studies of professional training from organizational psychology are useful for examining the influence of work (e.g., research) on professional identity development (e.g., being a science researcher). A longitudinal study of medical residents engaged in primary care, radiology, and surgery training tracks, found that residents attended to contextual factors, such as the work they were assigned and the professionals with whom they interacted, to develop their professional identities (Pratt et al., 2006). Similar to the science identity model from Carlone and Johnson (2007) and Hazari et al. (2010), Pratt and colleagues (2006) found that competence and recognition played an influential role in whether residents saw themselves as medical professionals. How residents described their professional identity was dependent on two interconnected cycles: work-learning and identity-learning. Residents experienced the **work-learning cycle** as they engaged in assigned work and made assessments of how that work aligned to their notions of work in their field (i.e., what a physician in primary care, radiology and surgery does and how they do it). As residents conducted their work they received informal feedback that informed changes in their perceptions of competence. When residents found that the content and process of their work did not align with their notion of work in their field or when they saw themselves as incompetent, they reconceptualized their professional identity on a continuum from being a medical student, a “baby” doctor, to eventually a fully-fledged physician as part of the **identity-learning cycle**.

Here, we aim to make two main contributions. First, we integrate science identity and professional identity theories to understand how science identity develops among ECRs during research training. We use the science identity model to conceptualize individual-level psychological factors such as recognition of self, performance-competence, and interest at play during ECRs’ science identity development. We use the professional identity model to elucidate aspects of research work that influence ECRs’ science identities, as well as how ECRs make sense of the factors that they perceive to affect their identity (Vough et al., 2020). Second, we synthesize findings from ECRs and return to the literature to identify relevant concepts and constructs that comprise and affect science identity development to generate a new model of science identity. Our model proffers fresh theoretical insights by accounting for the growth that can occur in science identity as individuals engage in research training experiences. Through two phases of research, we address the following research questions: (1) How do ECRs conceptualize their science identity and its potential for growth? (2) What aspects of ECRs themselves and their research experiences and environments influence whether and to what extent they identify as science researchers? and (3) How do ECRs make sense of the aspects of themselves and their research environments that influence their science identity?

## Methods

This investigation included two distinct phases of data collection and analysis. Both phases were deemed exempt by the University of Georgia’s Institutional Review Board (PROJECT00000870 and PROJECT00005063). All participants provided informed consent. In Phase 1, we surveyed undergraduates who had completed at least one term of life science research about whether and how their science identity changed because of their research experience(s). In Phase 2, we conducted semi-structured interviews with undergraduate, postbaccalaureate, and doctoral researchers about whether and how they identified as a scientist, including how their science identities were shaped by their research experiences. Together, these phases afforded broad insights about undergraduate researchers’ science identities (Phase 1) and deep insights about ECRs’ science identities across research training stages (Phase 2). We describe the data collection and analysis methods for each phase below.

### Phase 1

#### Data collection

Data were collected as part of a larger study of undergraduate research experiences during fall 2020, spring 2021, and fall 2021. Students (n=670) in this study were conducting life science research either as an intern in a faculty member’s research group or as part of a course at one of nine universities across the United States. Here we focus on their responses to the survey prompt, which they received at the end of the term: “Has this research experience made you feel more or less like a scientist? Please explain.” A total of 548 students responded to the prompt (82% response rate; Table 1). Their responses ranged from a few words to several sentences.

**Table 1.**
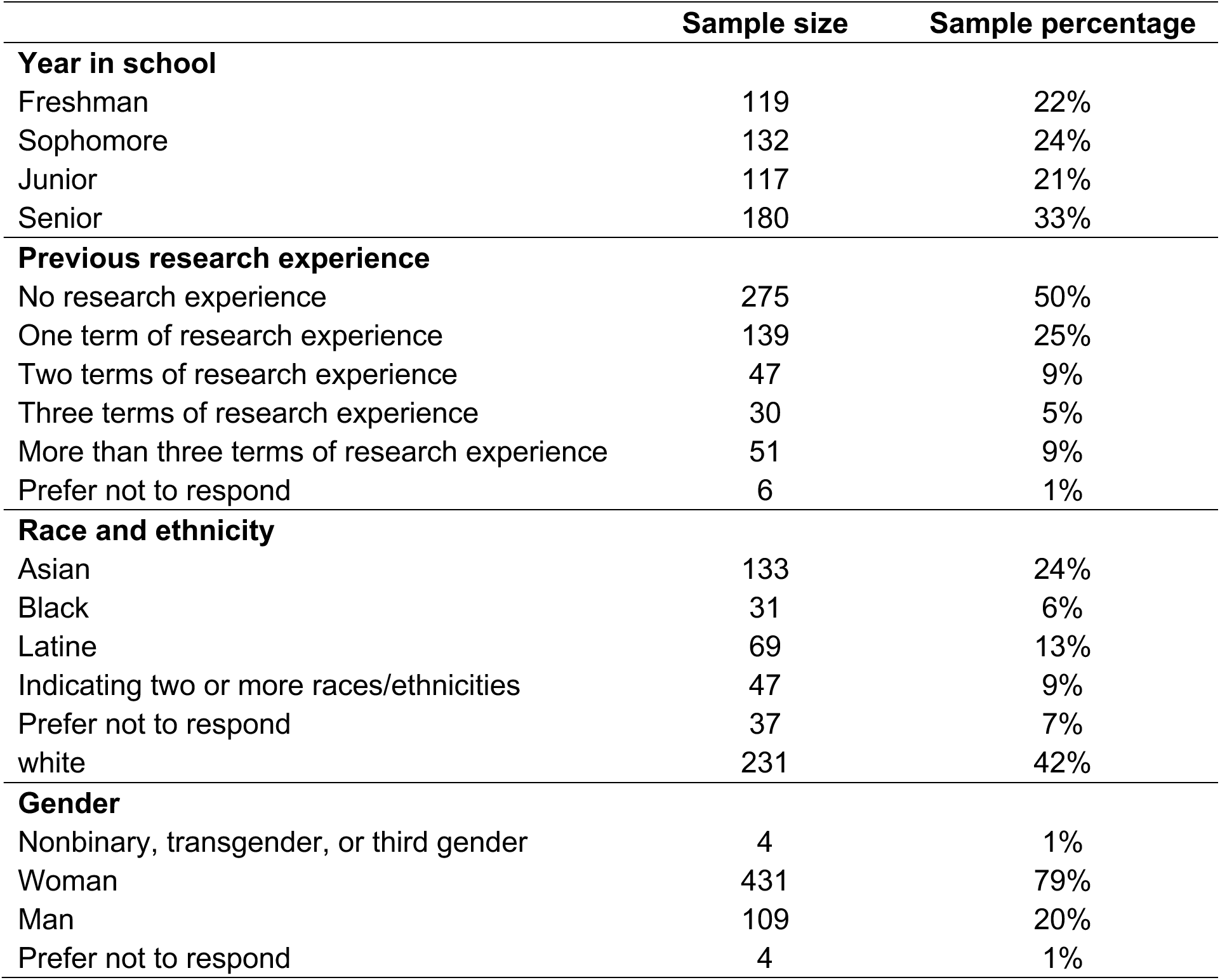
Summary of Phase 1 participant characteristics.

|  | Sample size | Sample percentage |
| --- | --- | --- |
| **Year in school** |  |  |
| Freshman | 119 | 22% |
| Sophomore | 132 | 24% |
| Junior | 117 | 21% |
| Senior | 180 | 33% |
| **Previous research experience** |  |  |
| No research experience | 275 | 50% |
| One term of research experience | 139 | 25% |
| Two terms of research experience | 47 | 9% |
| Three terms of research experience | 30 | 5% |
| More than three terms of research experience | 51 | 9% |
| Prefer not to respond | 6 | 1% |
| **Race and ethnicity** |  |  |
| Asian | 133 | 24% |
| Black | 31 | 6% |
| Latine | 69 | 13% |
| Indicating two or more races/ethnicities | 47 | 9% |
| Prefer not to respond | 37 | 7% |
| white | 231 | 42% |
| **Gender** |  |  |
| Nonbinary, transgender, or third gender | 4 | 1% |
| Woman | 431 | 79% |
| Man | 109 | 20% |
| Prefer not to respond | 4 | 1% |

#### Data analysis

We analyzed students’ responses qualitatively with the aim of understanding how their research experiences influenced their science identity. First, we reviewed and made analytic memos regarding our impressions of the data with the research question and ideas from the science identity model (i.e., performance-competence, recognition, and interest) and the professional identity model (i.e., work-learning and identity-learning cycles) in mind. Then we developed a qualitative codebook using a two-step process. First, we noted whether each participant perceived themselves to be more or less of a scientist at the end of their research experience, or if they were unsure of or unchanged in their perceptions. We then described what aspect(s) of their research experience informed their perceptions. We developed the codebook iteratively by proposing codes using an eclectic coding scheme. Specifically, we generated in vivo codes that use participants’ verbatim language in the code name, descriptive codes that use nouns to capture the topic of the data segment, and process codes or gerunds to summarize actions described in the data (Saldaña, 2016). We applied our initial codes to subsets of the data, and then met as a group to discuss the codes and associated data. During these discussions, we revised code definitions as needed and resolved any coding disagreements, producing the next iteration of the codebook. We applied the refined codebook to a new subset of data until the codebook stabilized. We then coded all data to consensus using the final stabilized codebook. At least two researchers applied the final codebook to all of the data and agreed on the application of the codes. At the end of this “first-cycle” coding process, we grouped the codes into larger, more abstract categories and themes while connecting them to the science identity and professional identity models guiding the study. First, we identified categories that summarized groups of related first-cycle codes. Then we grouped related categories into themes. The outputs of this analysis were further analyzed and synthesized with Phase 2 findings.

### Phase 2

#### Data Collection

To more deeply understand ECRs’ science identities, including how they evolved during research training, we conducted an interview study of undergraduate, postbaccalaureate, and doctoral researchers (i.e., ECRs) from several natural science disciplines (e.g., life sciences, geosciences, oceanography, environmental sciences). We recruited ECRs in spring 2022 using purposeful and snowball sampling. We identified potential participants by contacting research training program directors and principal investigators associated with science and technology centers funded by the National Science Foundation. We emailed study information to potential participants and made brief presentations about the study during research group meetings. We asked individuals interested in participating to complete a screening survey about their current level of research training, program or institutional affiliation, research focus, demographic information, and contact information (see Additional Information). A total of 57 individuals completed the screening survey. We invited 35 for interviews and 31 completed an interview: seven undergraduate researchers, seven post-baccalaureate researchers (postbacs), ten doctoral students who had not yet advanced to candidacy, and seven doctoral candidates (see Table 2). One interview was not analyzed further due to problems with recording quality, bringing the final sample to 30 participants. Participants were compensated with a $25 gift card.

**Table 2.** Summary of Phase 2 participant characteristics.

|  | Sample size | Sample percentage |
| --- | --- | --- |
| **Research methods primarily used** |  |  |
| Bench (including fieldwork) | 13 | 43% |
| Computational | 10 | 33% |
| Both | 7 | 23% |
| **Institution type** |  |  |
| Private doctoral university, very high research activity | 11 | 37% |
| Public doctoral university, very high research activity | 17 | 57% |
| Private baccalaureate | 1 | 3% |
| Public doctoral university | 1 | 3% |
| **Race and ethnicity** |  |  |
| Asian | 4 | 13% |
| Black | 2 | 7% |
| Latine | 5 | 17% |
| Indicating two or more races/ethnicities | 1 | 3% |
| Prefer not to disclose | 2 | 7% |
| white | 16 | 53% |
| **Gender** |  |  |
| Nonbinary, transgender, or third gender | 2 | 7% |
| Woman | 14 | 47% |
| Man | 14 | 47% |

We developed a 14-question interview protocol based on preliminary results of our Phase 1 analysis (see Additional Information) to query ECRs about their research experiences, their views of science researchers, their views of the work that science researchers do, and how they see themselves with respect to these views. We also inquired about how their views had changed, if at all. After the first few interviews, we added two questions to enhance elicitation from participants. Specifically, we asked participants to describe what, if any, work they did that they do not consider to be the work of a science researcher, and to explain how the culture of their lab group affects their science researcher identity, if at all. These questions helped us understand how ECRs perceived the nature of research work and the influence of their lab environment. The first author conducted all interviews over ZOOM™ using a semi-structured approach; interviews lasted ∼1 hour.

#### Data analysis

We had interview recordings transcribed by a professional transcription service (Rev.com) and we checked the transcripts for accuracy prior to analysis. We analyzed the interview data using a two-step process similar to our Phase 1 methods in which we developed a new codebook and engaged in coding to consensus. The Phase 2 dataset necessitated the use of versus codes as well as in vivo, descriptive, and process codes to make sense of the data. Versus codes use dichotomous terms to reflect tensions individuals exhibit within themselves (Saldaña, 2016), such as belonging versus being excluded. For the Phase 2 dataset, we conducted a more elaborate second-cycle coding process that involved pattern and axial coding of first-cycle codes that addressed our research question (Saldaña, 2016). We referred to existing literature sources to make sense of the ideas emerging from the analysis as needed. Through second-cycle coding, we again identified categories and themes that we then compared to the categories and themes from Phase 1. We refined the descriptions of our categories and themes to encompass findings from both phases. Descriptions of the emergent categories and themes along with example data are included in Additional Information File 3.

Quote data we present have been lightly edited for readability. Brackets indicate words added to enhance clarity. Ellipses represent words or phrases removed for conciseness. Phase 1 responses are indicated in text and Phase 2 ECRs are assigned pseudonyms.

#### Trustworthiness and positionality

We endeavored to conduct and present a trustworthy qualitative study. We provide descriptions of how we collected data and highlight key analytic decisions and processes used to arrive at the themes and categories we propose (Tracy, 2010). Throughout the course of our study, we wrote analytic memos and detailed each decision we made along with our rationales for these decisions (Charmaz, 2006). We engaged in forms triangulation by collecting and analyzing two distinct datasets and by coding data to consensus with multiple researchers (Krefting, 1991). During our analysis, we presented our findings to researchers unfamiliar with our data corpus. We also engaged in a form of member checking by presenting a summary of our findings at a research meeting where ECRs in the study could view and comment on our results (Krefting, 1991). Questions and comments from these sessions helped us to clarify the findings and implications of our study as well as enhance their trustworthiness.

Another component of trustworthiness is acknowledging our positionalities that may influence how we interpreted our data (Suddaby, 2006). As a research team, we represent perspectives of faculty, postdoctoral researchers, graduate students, and undergraduate researchers familiar with research training experiences in the natural sciences. We engaged in bracketing as a tool to recognize our positionalities. Bracketing helps researchers identify and reflect on how their own experiences of the topic they investigate may manifest during the research process and complements analytic memo writing as a form of reflexivity (Tufford & Newman, 2012). Specifically, we wrote responses to the questions: (1) How do I see my own identity as it relates to science and/or research? (2) In what ways, if any, am I like “an insider” to the ECRs in this study? (3) In what ways, if any, am I like “an outsider” to the ECRs in this study? (4) How could my own views and perspectives influence data analysis? A summary of researcher responses to the bracketing exercise are provided in Additional Information. By coding to consensus as well as clarifying and revising findings based on feedback from the member check and researchers unfamiliar with the data corpus, we strove to mitigate the effect of any individual researcher biases.

#### Limitations and considerations for transferability

Several aspects of our study have the potential to limit the transferability of our findings. First, our Phase 1 sample has a higher proportion of women than men. Second, ECRs in both phases volunteered to participate in studies of research training and our Phase 2 sample was comprised of ECRs who agreed to participate in a study regarding research experiences and researcher identity. Thus, our sample is likely to be enriched for individuals with a relatively robust science researcher identity, or who may otherwise feel comfortable discussing their science identities. We purposefully selected ECRs in Phase 2 to optimize the diversity of our sample in terms of amount of research experience, research type, race/ethnicity, and gender. Our Phase 2 sample is relatively representative of the national population of undergraduate and graduate students in terms of race, ethnicity, and gender (National Science Foundation, 2021). We did not prioritize participant selection by institution type. Most of the ECRs in Phase 2 are from institutions with very high research activity.

## Results and Discussion

Early career researchers (ECRs) in our study conceptualized science identity as having three forms: science student, science researcher, and career researcher. Their descriptions of these forms relied primarily on their views of the purpose of an individual’s daily work and the types of tasks typically executed by individuals at each level. ECRs who recognized themselves as “science students” tended to call themselves science majors or science graduate students and saw the purpose of their daily work as learning about research or learning to conduct research. ECRs who recognized themselves as “science researchers” tended to see their daily work as addressing research questions or testing hypotheses. No ECR in our study described themselves as a “career researcher” because they perceived the daily work of career researchers to be securing funding, setting big-picture research goals, and mentoring a team of researchers.

### Autonomy and epistemic involvement

ECRs’ recognition of themselves as science students or science researchers depended on the extent to which they perceived they had autonomy in their research. ECRs’ descriptions of “autonomy” resembled the notion of operational autonomy, or “the freedom, once a problem has been set, to attack it by means determined by oneself, within given resource constraints” (Bailyn, 1985, p. 134). ECRs who reported having operational autonomy described having freedom to carry out research tasks on their own, set their own hours, decide with whom they work, and, for some types of research, choose where they conducted their research. They also indicated that their research advisors or mentors were “hands off,” implying that they could decide how to execute their research. For instance, Shae talked about how her principal investigators (PIs) are available if she needs help, but she otherwise works independently:

> If I’m trying to run a specific experiment or work through this protocol, they’re not standing over my shoulder making sure that I’m doing everything properly. They’re hands off in the sense that they let us try things ourselves and if something is not working and we ask them for help, then they’ll step in.

Lisa conveyed that she came to understand that being a researcher means learning to navigate the operational autonomy that is inherent to her work:

> [As an undergraduate researcher] I was kind of helping someone else. Someone else was guiding me and providing help. [Now] I have to decide what I’m doing on a day-to-day basis. I get to decide what time I come into work and when I leave.

ECRs explained how they developed greater operational autonomy as they gained experience, as Harris stated:

> In your earlier days of grad school, you can’t choose who you are going to work with, especially if your PI assigns a certain mentor to your research. As a senior graduate student, I feel like I have more freedom in choosing who I want to work with.

All of the ECRs in our study indicated that their research decisions and directions were still guided and approved by their PIs, at least to some extent. Thus, none appeared to have strategic autonomy, or “the freedom to set one’s own research agenda” (Bailyn, 1985, p. 134). Rather ECRs described strategic autonomy as unique to being a career researcher, as articulated by Jane:

> I don’t have the freedom of creativity that I would think that a typical [career] science researcher would have in terms of making my own project. But I am a researcher in the sense that I am actively doing research and like contributing to projects…[I’m] just not at the top level yet.

Some ECRs acknowledged that they would feel more like a researcher once they experience full strategic autonomy, as Claire described:

> I think I will feel more like a science researcher when I get to the point of fully creating my own project and answering questions from scratch, because up to this point, I still need some guidance. I think that is really common in [starting ECRs]. I was never going to walk in and be like, “I want to do this.” Especially with my first project, it’s building off of others. I have this dataset and a lot of direction…when I get to the point where I can direct my own research, regardless of what it is, I think that’s when I’ll feel like a researcher.

For Claire, “directing” her own research (i.e., developing strategic autonomy) is the point at which she will consider herself a fully-fledged science researcher.

The more experienced researchers in our sample described how they were building towards full strategic autonomy by engaging in epistemic involvement, or taking intellectual responsibility for the research (Burgin et al., 2012). ECRs who felt they had limited epistemic involvement, such as being responsible for just “executing protocols” with little knowledge of the purpose of their work or limited involvement in data analysis, tended to see themselves more like a science student. ECRs who reported greater epistemic involvement, such as by formulating or refining research questions or directions and troubleshooting unexpected results, identified more as science researchers. To recognize oneself as a science researcher, ECRs described needing to contribute beyond “collecting data.” For instance, Skylar described how he did not feel like a researcher when he was working as a technician for a government agency before graduate school, revealing, “I just felt more like a gear in a larger machine…out there, collecting data and then handing off all that data to somebody else to analyze and draw conclusions from.” Justin echoed Skylar’s views, describing that he does not really feel like a researcher right now due to a lack of epistemic involvement:

> I do participate in research, but in terms of like the actual assay development and the thought processes behind this stuff, I’m not as involved as I was in my previous research [experiences] [in] undergrad. [There] I had full autonomy on my project. [Now] I basically just process samples and just get stuff done. I don’t have to think too deeply about what I do outside of how to be consistent and how to make sure things are in place so that people can do their jobs…I’m not involved in the decision making for the projects.

ECRs described additional ways they were epistemically involved. For instance, Levi shared that he saw himself as a researcher in one of his labs because he contributed intellectually to the troubleshooting process when experiments go awry:

> I do make suggestions, and sometimes those suggestions are right. One time I made a bunch of agar with a different, a new brand of agar, and nothing grew on it. I wrote everything down, but [my mentor] didn’t realize that I had to use the new brand since we were out of everything else. I was like, “Oh, this is probably the agar that’s the issue.” We did another test because of what I found out, and it was indeed the agar.

Levi continued to reflect on why this made him feel like a researcher “…research wise, I think I add my own ideas to what’s going on. I’m not just doing the program and doing the protocol, I’m definitely adding value to the research.” Other ECRs also described their epistemic involvement as evidenced by their contributions as Hazel expounded:

> [The research I do], it’s not totally independent. I do have supervision from the grad student I work under, but it’s like I’ve grown to be more independent in what I’m allowed to do. [Another undergraduate researcher and I] led a project where we came up ideas we would present to [to our graduate student mentor]. [Saying] like, “Hey, we wanna do this. Or we think that we should test this.” Then she would use her expertise [to] veto or [to] okay [our ideas].

Some ECRs described their epistemic involvement in terms of recognition, such as being an author on a paper, as Valerie explained:

> I’m finally gonna be a co-author on the paper. I made it, this is science. I feel like, since I’ve been able to contribute to things genuinely and contribute knowledge, I do feel like a researcher now. It took a while to get here.

ECRs in doctoral programs displayed epistemic involvement when designing chapters of their dissertations. Skylar expressed that he is trying to design his chapters in a way that addresses his own research interests and the interests of his PI, “[I am] just trying to find some link between, what my advisor wants me to do and what I like, to sort of dovetail and just sort of merge paths.” For ECRs, “merging” their own interests as well as their thoughts about the systems they investigate with their PI’s interests and thoughts supported their development of a science researcher identity.

Interestingly, ECRs exhibited varied reactions to being epistemically involved in their research. Justin stated that he feels “a relief being on this side of things” in his current role where he is not involved epistemically, although his work can get “monotonous sometimes.” Other ECRs, like Harris, find epistemic involvement, operational autonomy, and the potential for strategic autonomy appealing. He revealed:

> I guess one aspect of why I chose to go to grad school and took this career path is that it really allows me to be an independent person and to form my own projects that I’d like to work on. I get to set my own plans instead of someone else setting them for me.

Justin’s and Harris’s perceptions of epistemic involvement suggest that ECRs perceive and experience epistemic involvement in different ways, which is noteworthy because ECRs delineated the forms of science identity based on epistemic involvement and autonomy (operational and strategic).

### Individual factors

ECRs indicated three individual-level factors that affected whether they identified as science researchers: research and researcher conceptions, research skill perceptions, and career intentions. Each individual factor and how it shaped ECRs’ identification as a science researcher is described below.

#### Research and researcher conceptions

In describing whether they perceived themselves as science researchers, ECRs noted the qualifications they viewed as necessary to be a science researcher. A Phase 1 ECR indicated, “I used to think being a scientist meant that you had to be a genius who was always right. Now I understand that most scientists actually fail, but it is still valuable.” Other ECRs made statements like, “I feel like I’m not a real scientist until I get my PhD.” These conceptions of what it is to be a researcher influenced how ECRs saw their own science identity. These findings align with prior research showing that ECRs vary in their researcher conceptions in ways that affect their science identity development (Åkerlind, 2008; Atkins et al., 2020; Brew et al., 2016; Meyer et al., 2005; Vasquez-Salgado et al., 2023; Zuckerman & Lo, 2022).

ECRs also felt more or less like a scientist based on their conceptions of their research – specifically whether they viewed it as authentic or “real.” One ECR mentioned that their training experience “made me feel more like a scientist. I’m doing real research and that makes me feel super valid.” Several Phase 1 ECRs felt less like a scientist because they did not consider the work they did to be “real” research. For example, an ECR responded, “This [research experience] has made me feel less [like a scientist]. I do not really think of doing much with computers as ‘science.’” Another ECR reported, “My research experience was less scientific (for example, we did not work in a lab looking at cell cultures), so it is hard to say that it made me feel more like a scientist.” Phase 2 ECR, Phoebe, elaborated on this notion of authenticity. When she first started her computational research project, she was not aware “that computational research was even a realm of research.” She elaborated that she “never had the desire to learn code” before her research experience, but she was “glad” to have done so. Through Phoebe’s research training experience, her view of research broadened. Broadening her conception of research to include computational work helped Phoebe see herself as a science researcher.

#### Research skill perceptions

ECRs also saw themselves as science researchers based on their perceptions of their own research skills and accomplishments, in line with the identity element of performance-competence (Carlone & Johnson, 2007; Hazari et al., 2010). ECRs tended to describe their research skills as insufficient or sufficient, which hindered or supported their science identity. A Phase 1 ECR divulged that “this [experience] has been difficult in that much of the scientific reading is very difficult to understand, causing me to often feel incompetent and less like a scientist.” ECRs who considered their research skills to be sufficient articulated how this view helped them feel more autonomy in their work, which, in turn, led them to recognize themselves as science researchers. A Phase 1 ECR who viewed their own research skills as sufficient stated, “It made me feel more like a scientist because I learned a lot about lab techniques and equipment to the point in which I can execute protocols on my own or know how to ask for help if needed.” ECRs cited examples of having sufficient research skills as reasons why they saw themselves as a science researcher during the Phase 2 interviews. Max shared that, when he questioned his own identity as a science researcher, he reviewed past versions of his curriculum vitae (CV):

> I’ll just kind of look down on myself [and think] maybe you’re not good at this [research]. What I’ve done is I’ve edited my CV over the last few years…What I’ll do is, I’ll go look back at previous versions of my CV and look to my CV now. And it’s like, you have done a lot…Look at all this on my CV. I have done all this…I am a science researcher.

Max reminded himself that he possesses sufficient research skills and has experienced epistemic involvement in research as evidenced by presentations and publications. This practice reinforced his science identity in moments of doubt.

#### Career intentions

ECRs saw themselves more or less as science researchers based on their career intentions. Some Phase 1 ECRs reported that they did not see themselves as scientists now because they did not intend to become a scientist in their future careers. Other Phase 1 ECRs detailed how their research experience helped them clarify the type of research they wanted to conduct in the future as they pursued a research-related career. One ECR responded:

> I feel more like a scientist because this is the first time that I’ve done pure analytical/dry lab research and not experimental/wet lab research. I prefer dry lab over wet lab, as I feel like I understand it and enjoy it much more.

Phase 2 ECRs described refining their career intentions throughout the course of their research training experience, which affected how they recognized themselves as science researchers. The career intentions of ECRs varied and included research-related or research-adjacent careers in academic teaching, academic research, government, industry, medicine, and non-profit organizations. ECRs sometimes described possible selves, or an “individual’s ideas of what they might become, what they would like to become, and what they are afraid of becoming” (Markus & Nurius, 1986, p. 954) when discussing their career intentions. Possible selves are known to evolve during identity development broadly and during science identity development more specifically (Calabrese Barton et al., 2013; Dunkel & Anthis, 2001).

The possible selves ECRs described centered upon whether or not they saw themselves becoming an academic PI and what other types of research-related or research-adjacent careers they may enjoy. Only a handful of ECRs explained that they aspire to become academic PIs because they think they are well-suited to the job. Iris elaborated:

> I think I’d make a really good PI because I like coming up with ideas and ways to tackle problems, but I don’t love lab work. That’s not my favorite part of science. I’d much rather work on data analysis and interpretation where[as] I know for a lot of people it’s the other way around. And I like writing. So, it’s comforting knowing that what I’m really good at is what I’m going to have to spend more of my time doing if I stay in academia [as a PI].

Some ECRs who aspired to become an academic PI wanted to do so because they enjoy teaching and mentoring. Seth stated that up until recently his career intention was to do research in industry. However, “I’ve taken an interest to becoming a professor out of nowhere, I think because of the [course-based undergraduate research experience I’ve been involved in]. I’ve seen that I really do enjoy teaching.” Another ECR, Skylar, hopes to become a PI, in part, because he enjoys mentoring. He stated:

> I don’t see myself as like one of the “big name” researchers or like a researcher that would ever even really want to be that big name. I don’t wanna rewrite the *Principles of Ecology* or anything like that. I just wanna ask cool [research] questions and mentor students. I’m hopeful to work in academia as a professor of research with teaching duties…specifically like [at] an R2 public university.

Skylar spoke to a perception held by many ECRs that remaining in academia at a research university means excessive work hours with relatively low pay that may jeopardize their work-life balance. Emma wanted to pursue a career in industry because in academia she sees:

> “A lot of bureaucratic bullshit. [An academic researcher] doesn’t get paid as much [as a researcher in industry]…As much as I love teaching now, that’s not something I want to do as a job. Academia’s a big no…it’s the money, mainly, and it just seems so stressful. I do like mentoring, but I’d rather do a one-on-one [mentoring], rather than mentoring a whole lab.

Like Emma, other ECRs did not want to become an academic PI, in part, because they perceived the job to require them to mostly focus on issues of lab administration as well as securing funding for research, which would prevent them from engaging in hands-on research regularly. These and other negative perceptions of academia fed into ECRs’ feared selves, or what they do not desire becoming in the future. Talia expounded on what she does not want to become as she discussed her future career plans:

> Academia is so brutal…I don’t wanna be a postdoc who’s going through like multiple rounds of job application cycles. And let [academia] chew me up and spit me out. I’ve seen enough unhappy people in academia that if [academia] starts to make me unhappy, I’m out. I can see myself doing research in the future [but] I’m probably more interested in making sure that I am teaching, [doing] some sort of work with people in the future. It’s probably more important to me than, than making sure I still do research.

As Talia alluded to when she said “I’ve seen enough unhappy people in academia…,” career intentions and possible selves were often connected to the social interactions and group norms ECRs experienced during research training. Our findings underscore that the context of research training informs ECRs about what a future career as an academic PI may look like.

### Contextual factors

ECRs described how the nature of their research work, their social interactions during research, and the norms within their research groups or institutions influenced their science identity development. Each contextual factor and how it shaped ECRs’ identification as a science researcher is described below.

#### Nature of work

ECRs discussed how the topic of their research work and the process of conducting this work influenced whether they saw themselves as science researchers. ECRs who had experienced multiple types of research tended to speak most directly about how the nature of their work affected how they recognized themselves as science researchers. Mitchell who conducted fieldwork, bench research, and computational research reported:

> [In one of my research labs] I did fieldwork with them. I was like out in the field, like on my knees, digging up dirt, getting bit by a thousand mosquitoes, interacting with the things that I am studying. [My work for that project] has really enabled me to develop that relationship with my work and with the people in the lab. That experience, that definitely informs how I feel as a science researcher. I feel like it really genuinely gives me more authority on things because I have one more step of understanding how this [system] works because I’m out there looking at it.

For Mitchell doing “hands-on” work bolstered his science identity. Harris, who had experience conducting both bench and computational research described how the nature of work informed him of the type of researcher he wants to be:

> [As a computational researcher] I definitely have more independence in terms of when and where I want to do my work. If you’re doing bench work, your schedule is sometimes tied to how your cells grow and how your experiments turn out. One thing that I do admire about bench work is that you actually get to see things work in the real world…[In computational research], results only happen inside a computer. You can’t really visualize [your results] in the real world and actually see your successful results, I guess [as a computational researcher]. I feel like that is definitely the more rewarding part in terms of doing actual bench work. But in terms of the freedom and time, that time and space that I have, I feel like I just value that more.

The freedom Harris perceived to be inherent in computational research ultimately influenced his intention to become a computational researcher. These findings illustrate how the nature of work can influence how ECRs recognize themselves as science researchers.

#### Social interactions

ECRs spoke about the scope and quality of social interactions they experienced in research settings and how this functioned as recognition of their epistemic involvement in their research, in alignment with the recognition element of science identity (Carlone & Johnson, 2007; Hazari et al., 2010). Notably, ECRs perceived social interactions as positively or negatively influencing whether they saw themselves as science researchers. For example, Sutton conducted computational research and was involved in making tools to help onboard new members of the research team. Sutton viewed this work as important because it enabled research. During graduate school interviews, Sutton learned that not all researchers considered their work to demonstrate epistemic involvement and described this negative social interaction:

> One of the terms that people use that is derogatory in programming or [computer sciences] is [the term] “code monkey.” [It means] someone who just is given a task and [told] to go code it, without any creative input. [During my interview] I was asked, “Are you doing modeling work for them? Or are you just like a code monkey in this lab?”

Sutton interpreted the interviewer’s comments to mean that they were not a “real” science researcher in the eyes of the interviewer and they started to question this themself. After taking time to consider the interaction and its meaning, Sutton reaffirmed that they saw themself as a researcher because their current research work was producing scientific knowledge:

> It’s unfortunate, perhaps, to an extent [that this interaction happened to me], but it is something that I’m keeping in mind now. [I realized] other people may not respect [the work I’ve done] as much as the people in my lab, who are going to use [what I did] on a regular basis…Now, I’m working on what this person [in the graduate school interview] might consider to be science.

Sutton’s example highlights how epistemic involvement is the currency by which science researcher identity is recognized.

A lack of social interactions also influenced ECRs’ perceptions of their own epistemic involvement, which in turn influenced how they saw themselves as science researchers. Phase 2 ECR, Jane, explained elaborated: “I honestly barely get any interaction [in my research]…A lot of it’s solo work. It’s very, very hands off.” She shared that in her previous research in a different lab she experienced more social interactions and there she felt that she was recognized for epistemic involvement. “I knew everybody and knew exactly what my role was and where my place was and what I was good at and what people needed me for.” She shared that in her new lab because it’s her just “flying solo” she misses recognition of her epistemic involvement in the lab by her peers and that her science identity was languishing as a result. “I don’t really feel like anyone, no one really relies on me here yet. I’m trying to build that. To be like a go-to person for a certain thing.” Jane reported that she is actively seeking more collaborative projects in her current lab. She reported she would feel more like a science researcher when she experienced social interactions that reinforced her epistemic involvement.

ECRs identified how positive interactions with lab members and with people outside the lab helped them feel more like a science researcher by enhancing their perceptions of their epistemic involvement and autonomy in research. For example, Phase 2 ECR, Phoebe, explained how her PI helps her recognize herself as a science researcher:

> Our PI alone, they almost believe in us before we do. They give us the freedom to do these projects when we don’t even think we’re capable. They truly want us to succeed and are setting us up to succeed, and then get really excited about our success.

Like Phoebe, Phase 2 ECR Valerie, articulated that ongoing positive interactions with her PI supported her science researcher identity. Valerie described working closely every day with her PI and that the trust they built in their relationship enabled the development of her science researcher identity. She stated: “Taking me on as a master’s student, I got to essentially choose what I wanted to do for my master’s, which is not common at all.…Now I have my dream project that I helped make.” Valerie will soon begin a master’s project advised by her PI, which she helped to define (i.e., epistemic involvement) and she would have the freedom to carry it out (i.e., operational autonomy). Like social interactions, group norms were a contextual factor influencing ECR science identity.

#### Group norms

ECRs described norms – or understood, expected patterns behavior or standard ways of operating– that shaped their identification as a science researcher. ECRs referred to norms at multiple levels, including in their research groups, academic departments, research training programs, institutions, and disciplines. The norms inherent to these groups served as a source of information ECRs used to gauge the traits, skills, and values that were expected of science researchers in their context. ECRs cited examples of how their lab norms centered on inclusive practices, which helped them feel more like a researcher. Examples ranged from the subtle to explicit. One subtle norm was the daily practice of greeting all researchers who were working in the lab when they came in to do research. A few ECRs reported that this was not a norm consistent in all the labs they had worked in. They reported that exchanging daily greetings with their colleagues was a way they felt recognized as a researcher. Explicit norms were established through direct conversations and endorsements by the PI about inclusive practices like how to communicate more effectively in the research group when members of the team speak a variety of first languages.

ECRs also reported how the reputation of their institution, or how other researchers within the same context were typically regarded, could affect how they viewed their work and thus their science identity. Skylar shared that he recently completed his master’s research in a “humble” research group. Although he conducts similar research in his doctoral studies, the reputation of his current doctoral research group changed how he sees the “validity” of the research he does and helped him see himself more as a researcher:

> Now I’m in this lab where it’s like super flashy…my lab mates, they’re on NPR [National Public Radio]…[The notoriety of the lab] makes [research] feel even more serious and more high stakes…your lab group can definitely shape how you see yourself as a researcher.

ECRs indicated that observing other individuals, including ECRs and career researchers, affected what they thought was possible for themselves. These data supported the notion of ECR possible selves being influenced by contextual factors (Markus & Nurius, 1986). Some ECRs discussed modifying their career goals based on what they observed in their research groups and departments. Ezra stated:

> When I got here, I wanted to be the very best, I wanted to be the top scientist in the world. When I realized what that means and what that [lifestyle] is like, I [decided] I don’t want this in my life. I do want to be a science researcher, and there’s other ways I can do that. I do not strive to be someone who publishes in [a prestigious journal]…At some point in life, I wanted to be that person. But now it’s like, I can still be a science researcher, but I don’t want to be at the level of these people.

Ezra and other ECRs shared that observing the long work hours of other researchers, especially faculty, caused them to question if they would be happy or successful in that type of career.

Some ECRs discussed how seeing researchers who looked like themselves or held similar aspirations helped them feel like a research career was viable, and this supported their science identity development. Shae, who described wanting to start a family and pursue a research-related career, shared what it meant for her to see a graduate student have two children and be successful in research:

> When I was younger, and I even sometimes think about this now, [I wondered] how do women do this [pursue research and research-related careers] and have families or have a personal life? It seems like balancing and juggling all of that must be so difficult. Seeing a PhD student in my first lab have children was really influential for me because it was the first time that I was able to see that women are able to have families and children but still be very successful in their field.

For Shae, seeing women with children in a research lab informed her that being a science researcher and a mother is possible. Group norms, thus, can affect individual factors like career intentions.

### Making sense of influential factors

Overall, ECRs reported variation in the degree of challenge they experienced forming and sustaining a science researcher identity. Several ECRs explained that they recognized themselves as a science person for most of their lives and perceived recognizing themselves as a science researcher to be relatively straightforward. Other ECRs described how they made sense of individual and contextual factors as part of an ongoing process of identity development. Indeed, prior research indicates that sense-making plays a critical role in identity development (Vough et al., 2020). ECRs made sense of factors influencing their science identity using three main processes: perceived cohesion, attribution, and critical reflection. For instance, Lynn is an ECR from a marginalized racial background. She described having difficulty forming a science researcher identity when she started doing research:

> I just felt like I was throwing myself into a completely different persona…I wanted to do science, but I didn’t know how exactly I fit in science and how I could change the person that I felt like I was on the outside to be, I guess, that idea I had [of] a science researcher.

When asked what she felt like she had to change to be a science researcher, Lynn responded:

> I thought there was a certain way that you needed to dress. I tried to change that. I just had this idea that there were things that you’re supposed to do and ways that you’re supposed to act to make yourself a [science] researcher. I guess I tried to be more serious. Not really show my personality, only think about science.

In the interview, Lynn did not directly name racism in science as the reason she felt like she had to change herself to be a researcher, however, these data could be interpreted as cultural codeswitching in the face of racism (Cross et al., 2017; McCluney et al., 2021; McGee & Martin, 2011; Rincón & Hollis, 2020; Stanton et al., 2022).

Lynn appeared to make sense of her science identity using perceived cohesion, or by attending to “an attribute of individuals within a group that reflects an appraisal of their own relationship to that group” (Bollen & Hoyle, 1990, p. 482). She perceived her clothing and personality as not fitting those of the group, and thus she did not fit the group. These perceptions motivated Lynn to engage in community outreach at elementary schools and community colleges. She explained:

> [Outreach work is] important because I had no idea that being a scientist was something that was attainable. I had [an interest in] science as I was growing up, but I had never been told that that was something that I should pursue or had any example [of a science researcher in my life].

Lynn participated in outreach to show that people like her are science researchers. Additional ECRs discussed how they made sense of factors influencing their science identities.

Ezra identifies as a man from a marginalized racial background. He described that, at the start of his graduate school experience, he felt like he did not belong, “I just felt [like] you probably are not meant to be here, this is not for you” (i.e., perceived cohesion). He perceived himself as the source of the “problem” – that is, he attributed deficiencies within himself as the reason why he did not feel like he belonged or was truly a science researcher. After making friends and participating in diversity, equity, and inclusion-focused professional development, he viewed himself more as a science researcher because of how he attributed the source of the misfit. He reported that “it’s not that I don’t fit here. It’s that I was not allowed to be myself in this environment.” This statement indicated that he changed his causal explanation of misfit to one of circumstances or context rather than something about him (Graham, 2020; Weiner, 1985).

Furthermore, Ezra described that he came to this change in mindset by critically reflecting on his academic and research environments and how these environments were created without consideration for his own needs and identities. Ezra was critically reflective when he attributed the cause of the social inequity he experienced to system-rather than individual-level phenomena (Watts et al., 2011). Ezra described the development of critical reflection as a “change” in his “mindset” and in the language he used, which he found to be “very powerful” for seeing himself as a science researcher. Other ECRs engaged in critical reflection that was supportive of building and maintaining a science researcher identity, especially in challenging research environments.

### Emergent conceptual model of science researcher identity

In Figure 1, we present a conceptual model of science identity development in the context of research training. This model integrates the science identity model prominent in science education (Carlone & Johnson, 2007; Hazari et al., 2010) and Pratt’s model of professional identity (Pratt et al., 2006). This model also accounts for the influence of individual factors, contextual factors, as well as sense-making in forming and sustaining a science researcher identity. The questions embedded within each theme description summarize what ECRs described grappling with as they recognized themselves as science researchers.

**Figure 1.**
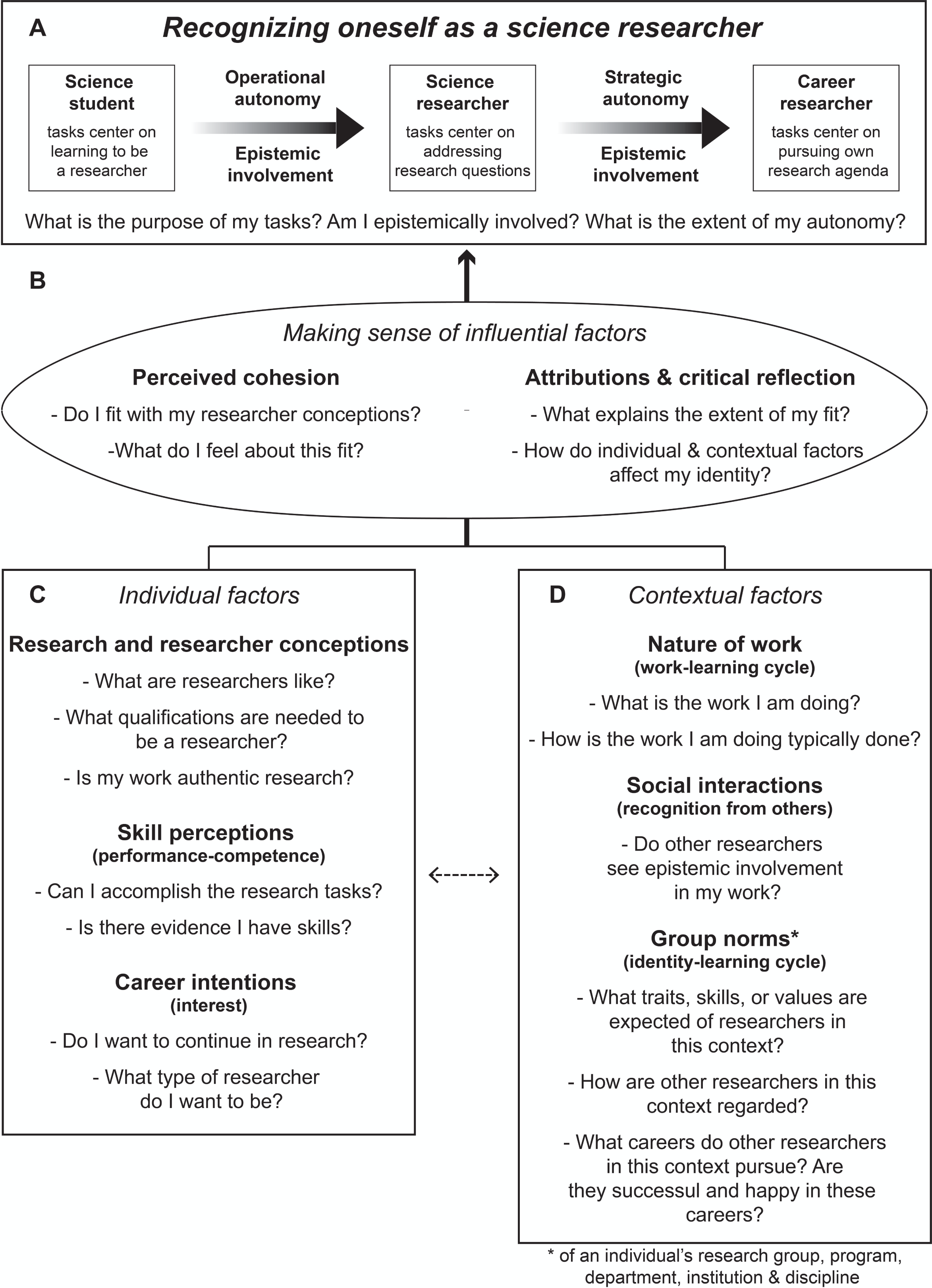
Conceptual model of science researcher identity development. The themes (italicized font) in our model are: recognizing oneself as a science researcher (A), making sense of influential factors (B), individual factors (C), and contextual factors (D). Each theme is composed of categories (bolded font) and questions (regular font) that ECRs contend with as they form and further develop their science researcher identity. Parentheses indicate instantiations from the models guiding this study: science identity (Carlone & Johnson, 2007; Hazari et al., 2010) and Pratt’s model of professional identity (Pratt et al., 2006). Recognizing oneself as a science researcher is bolded because it is the heart of our model. Shaded arrows represent increasing autonomy and epistemic involvement that differentiate the forms of science identity: science student, science researcher, career researcher. Themes, categories, and questions and arrows between the dimensions of the model are further described in the Results and Discussion.

- “Recognizing oneself as a science researcher” (Figure 1A) elaborates Carlone & Johnson’s science identity model (Carlone & Johnson, 2007) by delineating forms of science identity (science student, science researcher, career researcher) based on the focus of the work, level of epistemic involvement, and extent of operational vs. strategic autonomy (Bailyn, 1985). To place themselves on this developmental continuum, ECRs considered: What is the purpose of the tasks I execute during research? Am I epistemically involved in my research? What is the extent of my autonomy?
- “Making sense of influential factors” (Figure 1B), represented as a lens, shows that ECRs perceived cohesion (Bollen & Hoyle, 1990), made attributions (Graham, 2020), and engaged in critical reflection (Watts et al., 2011) to make sense of individual and contextual factors, ultimately identifying more or less as a science researcher. During their sense-making, ECRs considered: Do I fit with my researcher conceptions? What do I feel about this fit? What explains the extent of my fit? How do my individual and contextual factors affect my identity?
- “Individual factors” (Figure 1C) include research and researcher conceptions, skill perceptions, and career intentions. Research and researcher conceptions are novel to this model, while skill perceptions reflect the notion of performance-competence and career intentions reflect the notion of interest from the science identity model and possible selves (Carlone & Johnson, 2007; Hazari et al., 2010, 2013; Markus & Nurius, 1986). In considering whether they were science researchers, ECRs asked: What are researchers like? What qualifications are needed to be a researcher? Is my work authentic research? Can I accomplish the research tasks? Is there evidence I have skills? Do I want to continue in research? What type of researcher do I want to be?
- “Contextual factors” (Figure 1D) include ECRs’ perceptions of the nature of their work, their social interactions, and group norms. “Nature of work” and “group norms” are instantiations of the work-learning and identity-learning cycle from Pratt’s (2006) model of professional identity, respectively. “Social interactions” are instantiations of recognition from others from Carlone & Johnson’s (2007) science identity model. In considering whether they were science researchers, ECRs asked: What is the work I am doing? How is the work I am doing typically done? Do other researchers see epistemic involvement in my work? What traits, skills, or values are expected of researchers in this context? How are other researchers in this context regarded? What careers do other researchers in this context pursue? Are they successful and happy in these careers?

### Interactions between themes

ECRs’ remarks revealed that their individual factors (research and researcher conceptions, skill perceptions, and career intentions) were influenced by their contextual factors (nature of work, social interactions, and group norms) and vice versa. In addition, the interactions between these factors fed into how ECRs made sense and recognized themselves as science researchers. A Phase 1 ECR indicated:

> I definitely feel more like a scientist after this research experience. This was my first experience taking the lead of my own project. My PI really pushed me to make decisions on what data to collect, how to collect it, etc. Usually, as an undergrad, I don’t take ownership of projects and make decisions, but follow out a set guideline decided by the PI, postdoc, or grad student, with little to no insight as to why or how the guidelines were decided. By being the sole decision maker, with only very light guidance from my PI, I learned so much more about the pros and cons of each decision I made. Additionally, as the one writing the manuscript, I learned the why of my experiment very early on and have a really good grasp on the importance and translation of my project and how it fits into the existing literature.

This ECR reflected on how the nature of their work (contextual) offered them room for epistemic involvement and operational autonomy and that their social interactions and group norms (contextual) affected their skill perceptions (individual). Similarly, Shae began to recognize her own science identity by gaining awareness of what it means to be a science researcher. She described learning what is expected of a science researcher by “participating in a lot of the rituals of the lab, like going to lab meetings, and [having] one-on-one meetings with the PI [contextual]. By having those experiences, I started to learn this is what it looks like to be a science researcher.” Shae’s understanding of what it “looks like” to be a researcher within a particular lab group provided her a point of reference to assess her own identity.

Other ECRs, like Brian, reflected on how individual factors, such as research and researcher conceptions, related with contextual factors to shape how they ultimately saw their science identity and its evolution. As an undergraduate, Brian sometimes considered his research skills to be insufficient (individual) and, at that time, he only considered faculty to be researchers (individual): “If you had asked me [then] if a grad student was a science researcher, I might have said no. But for a while now, I have definitely thought that grad students are science researchers.” Brian then explained that the norms of graduate students carrying out the work (contextual) prompted him to reconsider his conceptions of who is a researcher (individual):

> In undergrad, I sort of imagined that professors were the ones doing most of the work behind research. But [now] as far as I can tell, it’s almost all grad student labor. … I’ve definitely shifted in my thinking of who is doing the science.

Brian now considers himself and graduate students to be science researchers.

A final example of how individual and contextual factors interacted to influence ECR science identity comes from Alex. Alex conducted research that was somewhat outside the purview of most members of their research group. They sometimes do not see themselves as a researcher because they wonder if the work they are doing is research or not. That is, their research and researcher conceptions sometimes do not align with the nature of their work. Alex reflected on what helps them see themselves as a researcher.

> The projects I’ve gotten, the goals have been concrete. For my current project [my PI told me a specific goal] and said here are examples that have done this. But then [my PI] puts a lot of trust in me to figure out how to do it…I’ve been able to change whatever I want other than the goal of the project itself. I think that’s been fulfilling in itself because I’ve been able to have agency [in my work]. My PI also makes sure everyone [in the lab] knows why the work [I do] is important. Cuz when you’re [doing the type of work I do] it’s hard to tell how people [in the lab] will actually use it. During lab meetings, the PI will be like, oh this is one valuable research use of [my work]. They even give specific examples to this grad student [who] could use it on this data they’re currently collecting. I think that’s been the most fulfilling part more than just the freedom in some ways.

The autonomy and epistemic involvement within Alex’s research work (nature of the work; contextual) helps them recognize themselves as a science researcher. Alex implies that their conceptions of research and researchers (individual) centers on intellectual freedom to pursue a research question or goal by means they determine for themselves. Furthermore, when their PI talks about their work in lab meeting—a setting in which both social interactions happen and group norms can be established, maintained, or dismantled (contextual)—it reifies Alex’s perceptions that the work they do is the work of a science researcher and, in turn, supports their science researcher identity.

Connections between themes in the model are reflected in the arrows in Figure 1. The arrows in 1A are shaded with a gradient to indicate increasing epistemic involvement and autonomy (operational and strategic) as ECRs transition among identities. The arrows from 1C and 1D through 1B to 1A illustrate that individual and contextual factors are fodder for ECR sense-making through which they ultimately place themselves along science researcher identity continuum. The dashed arrow between 1C and 1D denotes that individual and contextual factors interact bidirectionally, each shaping the other.

### Implications of model

Our work offers novel insights into how ECRs come to see themselves as science researchers. Our conceptual model integrates science identity and professional identity models in a way that is useful for explaining the transition from being a consumer of science (i.e., a science student) to a producer of science (i.e., a science researcher) (Carlone & Johnson, 2007; Hazari et al., 2010; Pratt et al., 2006). Our model also theorizes key features of research tasks that appear to influence whether ECRs identify more strongly as a science student or as a science researcher – namely, the extent to which ECRs have epistemic involvement in and operational autonomy over their research. These relationships should be tested in future research to understand the ways in which epistemic involvement and operational autonomy influence science researcher identity development. For instance, is epistemic involvement for any aspect of the research sufficient for ECRs to identify as researchers? Or do ECRs need to be intellectually responsible for posing research questions or setting research directions to transition from identifying as a science student to a science researcher?

Our model elaborates on the individual-level factors that influence science identity development by adding research and researcher conceptions to existing identity constructs of performance-competence and interest (Carlone & Johnson, 2007; Hazari et al., 2010). In addition to weighing whether they were interested in, capable of, and recognized for their scientific work, ECRs in our study considered whether their work was “real” research and how their views of themselves compared to their views of what a researcher is. Importantly, ECRs noted that their research and researcher conceptions changed as they engaged in one or more research training experiences. In other words, their “standard” for identifying as a science researcher evolved as they gained more and different research experience. Prior investigations of science identity development among undergraduate researchers have emphasized belonging in a community of scientists, but have not framed belonging in a way that accounts for dynamic views of the reference community. Our findings suggest that it may be important to account for this dynamicity in studying identity development among ECRs. ECRs’ changing research and researcher conceptions may be one reason that identity measures have fallen short of passing tests of measurement invariance over time (Hess et al., accepted).

Our model connects individual-level identity constructs (performance-competence, interest) and previously identified contextual factors (recognition from others) with other contextual factors that are known to be at play in professional identity formation (Pratt et al., 2006). Specifically, ECRs in our study described shifts in their thinking about what research is and how research gets done depending on the types of research they were doing (i.e., work-learning cycles). In addition, ECRs weighed norms regarding researchers and careers in their research environments as they assessed whether they identified more or less as a science researcher (i.e., identity-learning cycles). By connecting self-perceptions and contextual perceptions with science identity development, our model posits novel relationships that can be tested in future research. For instance, ECRs who have researcher role models in their environments, meaning people doing similar types of work who share their traits, values, or career aspirations, may be more likely to identify as a science researcher. This hypothesis could be tested by examining whether and to what extent ECRs who report having research role models also identify as science researchers (Atkins et al., 2020; Haeger & Fresquez, 2016). At least some research indicates that role models can improve motivation and persistence of individuals from marginalized or minoritized groups in STEM (Atkins et al., 2020; Hernandez, Bloodhart, et al., 2018; Hernandez et al., 2020). However, the influence of role models in the day-to-day research work environment has yet to be fully explored. Research role models may be especially important for ECRs who already strongly identify with the scientific community, but are less firm on their identification with their research community.

The connection between science identity development and work-learning and identity-learning cycles also raises questions of how identity development should be studied. A recent, longitudinal study of Latine undergraduates involved in research training demonstrated the dynamic nature of science identity. Over the 18 months of their research training, students exhibited different patterns in their science identities: consistent over time (either identifying as a scientist or not), increasing quickly, increasing gradually, or decreasing over time (Vasquez-Salgado et al., 2023). Likewise, work- and identity-learning cycles are inherently dynamic and have been studied longitudinally to capture change over time (Pratt et al., 2006). Yet, science identity as well as work- and identity-learning cycles are not typically studied in a way that can fully capture the potential extent of their dynamicity.

Investigations of ECR engagement in research over time and in context is needed to fully reveal the shifting and responsive nature of identity development. Research approaches such as experience sampling or ecological momentary assessment may prove useful for gaining novel insights into how ECRs come to identify first as science researchers and then as career researchers, including the roles of individual, contextual, and sensemaking factors in this process (Gabriel et al., 2019; Shiffman et al., 2008; Stone & Shiffman, 1994). Such approaches offer the potential for both ecological validity of the findings because data would be collected in the research training context, and understanding of the dynamicity of identity development because data would be collected over time. For instance, at the start of their training experience, ECRs may not yet fully recognize themselves as science researchers because they are still in the process of developing their researcher conceptions, skill perceptions, and knowledge of the nature of work, and they may have limited epistemic involvement and autonomy in their research. As they gain experience, they may grow in their skill perceptions and, as a result, take initiative toward operational autonomy and epistemic involvement, ultimately adopting more of a science researcher identity. Alternatively, research mentors or supervisors may afford ECRs more operational autonomy and epistemic involvement, which ECRs then perceive as recognition of their capabilities and a sign that they are becoming a science researcher. Longitudinal, concurrent study of these variables in the context of research training could reveal these processes and patterns.

Finally, our findings revealed that ECRs used three overarching processes to make sense of individual and contextual factors in identifying more or less as a science researcher: perceived fit between themselves and their conceptions of researchers (i.e., perceived cohesion); attribution of misfit to themselves or external factors (i.e., attribution); and recognition of systemic inequities as a source of misfit (i.e., critical reflection). These processes functioned as lenses through which ECRs made judgments about whether they were a science researcher, or not. These findings open multiple avenues for future investigation. First, the fact that ECRs attributed misfit to themselves or to external factors raises questions of whether attribution retraining – so called wise interventions – could be used to promote ECRs’ science identity development (Casad et al., 2018; Walton, 2014; Walton & Wilson, 2018). Similarly, interventions to build ECRs’ critical reflection skills may help them recognize the power they have to embrace a science researcher identity and reject contextual factors that present identity threats (Moll et al., 1992; van Veelen et al., 2019; Yosso, 2005).

If future research supports our model, there are likely to be practical steps research advisors and training programs can take to afford opportunities for ECRs to reflect on and grow in their science researcher identity. First, advisors can consider the extent to which they provide opportunities and support for ECRs to take on greater epistemic involvement and operational autonomy in their research tasks, especially as they gain experience. Second, advisors and programs can work to ensure ECRs have opportunities to interact with diverse groups of researchers. Connecting with affinity groups locally and nationally, such as through professional societies and at conferences, could broaden ECR research and researcher conceptions. Web resources, such as Scientist Spotlights (Schinske et al., 2016) and affinity-specific social media handles (e.g., @BlackinMarineScience, @DisabledinSTEM) may also be useful for expanding ECRs’ conceptions of who science researchers are. Third, advisors and programs can help ECRs to document, reflect on, and recognize their skill growth through the use of individual development plans and competency-based assessments that are revisited annually to reveal change over time (Chang & Saw, 2021; Kuniyoshi, 2021; Verderame et al., 2018). Programs could also structure learning about the historical and sociocultural issues at play within the science research community to support ECRs’ development of critical reflection, which, in turn, may strengthen their science researcher identity (Camacho et al., 2021; Heberle et al., 2020; Vasquez-Salgado et al., 2023; Watts et al., 2011).

## Conclusion

The goals of our study were to characterize how science identity is conceptualized by ECRs as well as how it develops during research training, how individual and contextual factors influence science identity development, and how ECRs make sense of the factors that affect their own science identity. We generated a conceptual model that accounts for our findings and joins and elaborates upon existing models and constructs regarding identity development. Our results enrich our understanding of how science identity develops during research training experiences. Findings from our study are particularly helpful in understanding the nuances of how ECRs with some preexisting research experience recognize themselves as science researchers and what factors affect their self-assessments of science identity. Our results offer theoretical insights to pursue in future research and suggestions to help bolster science identity development within ECRs. Supporting development of science identity within ECRs is likely to help individuals persist within training experiences and ultimately pursue science and research-related careers.

## List of Abbreviations

ECR: early career researcher
PI: principal investigator
Postbac: postbaccalaureate researcher

## Declarations

## Availability of data and materials

Interview questions used in this study are available in Additional Information. Data generated and analyzed may be available from the corresponding author upon reasonable request.

## Authors’ contributions

MAP and ELD designed the study. MAP collected data. MAP, CJZ, JMI, OAE, and ELD analyzed data. MAP and ELD collaborated on manuscript writing. ELD obtained funding for the study. All authors read and approved the final manuscript.

## Acknowledgements

We thank our participants for sharing their time and perspectives. We also thank program coordinators and principal investigators who facilitated data collection for this study. We appreciate the Social Psychology of Research Experiences & Education lab and the Center for Chemical Currencies of a Microbial Planet for feedback on emergent results. Lastly, we thank members of the Biology Education Research Group at the University of Georgia for helpful feedback on a previous version of this manuscript.

## References

Adedokun, O. A., Bessenbacher, A. B., Parker, L. C., Kirkham, L. L., & Burgess, W. D. (2013). Research skills and STEM undergraduate research students’ aspirations for research careers: Mediating effects of research self-efficacy. Journal of Research in Science Teaching, 50(8), 940–951. https://doi.org/10.1002/tea.21102

Aikens, M. L., Sadselia, S., Watkins, K., Evans, M., Eby, L. T., & Dolan, E. L. (2016). A social capital perspective on the mentoring of undergraduate life science researchers: An empirical study of undergraduate–postgraduate–faculty triads. CBE—Life Sciences Education, 15(2), ar16. https://doi.org/10.1187/cbe.15-10-0208

Åkerlind, G. S. (2008). An academic perspective on research and being a researcher: An integration of the literature. Studies in Higher Education, 33(1), 17–31. https://doi.org/10.1080/03075070701794775

Alexander, J. M., Johnson, K. E., & Kelley, K. (2012). Longitudinal analysis of the relations between opportunities to learn about science and the development of interests related to science. Science Education, 96(5), 763–786. https://doi.org/10.1002/sce.21018

Alston, G. D., Guy, B. S., & Campbell, C. D. (2017). Ready for the professoriate? The influence of mentoring on career development for Black male graduate students in STEM. Journal of African American Males in Education (JAAME), 8(1), 45–66.

Ambrosino, C. M., & Rivera, M. A. J. (2022). A longitudinal analysis of developing marine science identity in a place-based, undergraduate research experience. International Journal of STEM Education, 9(1), 70. https://doi.org/10.1186/s40594-022-00386-4

Atkins, K., Dougan, B. M., Dromgold-Sermen, M. S., Potter, H., Sathy, V., & Panter, A. T. (2020). “Looking at myself in the future”: How mentoring shapes scientific identity for STEM students from underrepresented groups. International Journal of STEM Education, 7(1), 42. https://doi.org/10.1186/s40594-020-00242-3

Bailyn, L. (1985). Autonomy in the industrial R&D lab. Human Resource Management, 24(2), 129–146. https://doi.org/10.1002/hrm.3930240204

Bollen, K. A., & Hoyle, R. H. (1990). Perceived cohesion: A conceptual and empirical examination. Social Forces, 69(2), 479–504. https://doi.org/10.2307/2579670

Brew, A., Boud, D., Namgung, S. U., Lucas, L., & Crawford, K. (2016). Research productivity and academics’ conceptions of research. Higher Education, 71(5), 681–697. https://doi.org/10.1007/s10734-015-9930-6

Burgin, S. R., Sadler, T. D., & Koroly, M. J. (2012). High school student participation in scientific research apprenticeships: Variation in and relationships among student experiences and outcomes. Research in Science Education, 42(3), 439–467. https://doi.org/10.1007/s11165-010-9205-2

Calabrese Barton, A., Kang, H., Tan, E., O’Neill, T. B., Bautista-Guerra, J., & Brecklin, C. (2013). Crafting a future in science: Tracing middle school girls’ identity work over time and space. American Educational Research Journal, 50(1), 37–75. https://doi.org/10.3102/0002831212458142

Camacho, T. C., Vasquez-Salgado, Y., Chavira, G., Boyns, D., Appelrouth, S., Saetermoe, C., & Khachikian, C. (2021). Science identity among Latinx students in the biomedical sciences: The role of a critical race theory–informed undergraduate research experience. CBE—Life Sciences Education, 20(2), ar23.

Carlone, H. B., & Johnson, A. (2007). Understanding the science experiences of successful women of color: Science identity as an analytic lens. Journal of Research in Science Teaching, 44(8), 1187–1218. https://doi.org/10.1002/tea.20237

Casad, B. J., Oyler, D. L., Sullivan, E. T., McClellan, E. M., Tierney, D. N., Anderson, D. A., Greeley, P. A., Fague, M. A., & Flammang, B. J. (2018). Wise psychological interventions to improve gender and racial equality in STEM. Group Processes & Intergroup Relations, 21(5), 767–787. https://doi.org/10.1177/1368430218767034

Chang, C. N., & Saw, G. K. (2021). Individual development plan, mentoring support, and career optimism among STEM graduate students during the COVID-19 pandemic. American Educational Research Association (AERA) Annual Meeting.

Charmaz, K. (2006). Constructing grounded theory: A practical guide through qualitative analysis. Sage Publications.

Chemers, M. M., Zurbriggen, E. L., Syed, M., Goza, B. K., & Bearman, S. (2011). The role of efficacy and identity in science career commitment among underrepresented minority students. Journal of Social Issues, 67(3), 469–491. https://doi.org/10.1111/j.1540-4560.2011.01710.x

Cribbs, J. D., Hazari, Z., Sonnert, G., & Sadler, P. M. (2015). Establishing an explanatory model for mathematics identity. Child Development, 86(4), 1048–1062.

Cross, W. E., Seaton, E., Yip, T., Lee, R. M., Rivas, D., Gee, G. C., Roth, W., & Ngo, B. (2017). Identity work: Enactment of racial-ethnic identity in everyday life. Identity, 17(1), 1–12. https://doi.org/10.1080/15283488.2016.1268535

Dou, R., Hazari, Z., Dabney, K., Sonnert, G., & Sadler, P. (2019). Early informal STEM experiences and STEM identity: The importance of talking science. Science Education, 103(3), 623–637. https://doi.org/10.1002/sce.21499

Dunkel, C. S., & Anthis, K. S. (2001). The role of possible selves in identity formation: A short-term longitudinal study. Journal of Adolescence, 24(6), 765–776. https://doi.org/10.1006/jado.2001.0433

Estrada, M., Hernandez, P. R., & Schultz, P. W. (2018). A longitudinal study of how quality mentorship and research experience integrate underrepresented minorities into STEM careers. CBE—Life Sciences Education, 17(1), ar9.

Estrada, M., Woodcock, A., Hernandez, P. R., & Schultz, P. (2011). Toward a model of social influence that explains minority student integration into the scientific community. Journal of Educational Psychology, 206–222.

Frantz, K. J., Demetrikopoulos, M. K., Britner, S. L., Carruth, L. L., Williams, B. A., Pecore, J. L., DeHaan, R. L., & Goode, C. T. (2017). A comparison of internal dispositions and career trajectories after collaborative versus apprenticed research experiences for undergraduates. CBE—Life Sciences Education, 16(1), ar1.

Gabriel, A. S., Podsakoff, N. P., Beal, D. J., Scott, B. A., Sonnentag, S., Trougakos, J. P., & Butts, M. M. (2019). Experience sampling methods: A discussion of critical trends and considerations for scholarly advancement. Organizational Research Methods, 22(4), 969–1006. https://doi.org/10.1177/1094428118802626

Gentile, J., Brenner, K., & Stephens, A. (2017). Undergraduate research experiences for STEM students: Successes, challenges, and opportunities. National Academies of Sciences, Engineering, and Medicine.

Godwin, A., Potvin, G., Hazari, Z., & Lock, R. (2016). Identity, Critical Agency, and Engineering: An Affective Model for Predicting Engineering as a Career Choice. Journal of Engineering Education, 105(2), 312–340. https://doi.org/10.1002/jee.20118

Goodwin, E. C., Cary, J. R., & Shortlidge, E. E. (2022). Not the same CURE: Student experiences in course-based undergraduate research experiences vary by graduate teaching assistant. PloS One, 17(9), e0275313.

Graham, S. (2020). An attributional theory of motivation. Contemporary Educational Psychology, 61, 101861. https://doi.org/10.1016/j.cedpsych.2020.101861

Haeger, H., & Fresquez, C. (2016). Mentoring for inclusion: The impact of mentoring on undergraduate researchers in the sciences. CBE—Life Sciences Education, 15(3), ar36. https://doi.org/10.1187/cbe.16-01-0016

Harackiewicz, J. M., Smith, J. L., & Priniski, S. J. (2016). Interest matters: The importance of promoting interest in education. Policy Insights from the Behavioral and Brain Sciences, 3(2), 220–227. https://doi.org/10.1177/2372732216655542

Hazari, Z., Chari, D., Potvin, G., & Brewe, E. (2020). The context dependence of physics identity: Examining the role of performance/competence, recognition, interest, and sense of belonging for lower and upper female physics undergraduates. Journal of Research in Science Teaching, 57(10), 1583–1607. https://doi.org/10.1002/tea.21644

Hazari, Z., Sadler, P. M., & Sonnert, G. (2013). The science identity of college students: Exploring the intersection of gender, race, and ethnicity. Journal of College Science Teaching, 42(5), 82–91. JSTOR.

Hazari, Z., Sonnert, G., Sadler, P. M., & Shanahan, M.-C. (2010). Connecting high school physics experiences, outcome expectations, physics identity, and physics career choice: A gender study. Journal of Research in Science Teaching, 47(8), 978–1003. https://doi.org/10.1002/tea.20363

Heberle, A. E., Rapa, L. J., & Farago, F. (2020). Critical consciousness in children and adolescents: A systematic review, critical assessment, and recommendations for future research. Psychological Bulletin, 146, 525–551. https://doi.org/10.1037/bul0000230

Hernandez, P. R., Adams, A. S., Barnes, R. T., Bloodhart, B., Burt, M., Clinton, S. M., Du, W., Henderson, H., Pollack, I., & Fischer, E. V. (2020). Inspiration, inoculation, and introductions are all critical to successful mentorship for undergraduate women pursuing geoscience careers. Communications Earth & Environment, 1(1), 7. https://doi.org/10.1038/s43247-020-0005-y

Hernandez, P. R., Bloodhart, B., Adams, A. S., Barnes, R. T., Burt, M., Clinton, S. M., Du, W., Godfrey, E., Henderson, H., Pollack, I. B., & Fischer, E. V. (2018). Role modeling is a viable retention strategy for undergraduate women in the geosciences. Geosphere, 14(6), 2585–2593. https://doi.org/10.1130/GES01659.1

Hernandez, P. R., Hopkins, P. D., Masters, K., Holland, L., Mei, B. M., Richards-Babb, M., Quedado, K., & Shook, N. J. (2018). Student integration into STEM careers and culture: A longitudinal examination of summer faculty mentors and project ownership. CBE—Life Sciences Education, 17(3), ar50. https://doi.org/10.1187/cbe.18-02-0022

Hess, R. A., Erickson, O. A., Cole, R. B., Isaacs, J. M., Alvarez-Clare, S., Arnold, J., … & Dolan, E. L. (2022). Virtually the same? Evaluating the effectiveness of remote undergraduate research experiences. bioRxiv, 2022-01. Accepted at CBE-Life Sciences Education

Hidi, S., & Renninger, K. A. (2006). The four-phase model of interest development. Educational Psychologist, 41(2), 111–127. https://doi.org/10.1207/s15326985ep4102_4

Kim, A. Y., & Sinatra, G. M. (2018). Science identity development: An interactionist approach. International Journal of STEM Education, 5(1), 51. https://doi.org/10.1186/s40594-018-0149-9

Krefting, L. (1991). Rigor in qualitative research: The assessment of trustworthiness. The American Journal of Occupational Therapy, 45(3), 214–222.

Kuniyoshi, C. Y. (2021). Individual development plans, your strengths, your career, and your professional identity. 2929 SACNAS The National Diversity in STEM Virtual Conference.

Limeri, L. B., Asif, M. Z., Bridges, B. H., Esparza, D., Tuma, T. T., Sanders, D., Morrison, A. J., Rao, P., Harsh, J. A., & Maltese, A. V. (2019). “Where’s my mentor?!” Characterizing negative mentoring experiences in undergraduate life science research. CBE—Life Sciences Education, 18(4), ar61.

Mahadeo, J., Hazari, Z., & Potvin, G. (2020). Developing a computing identity framework: Understanding computer science and information technology career choice. ACM Trans. Comput. Educ., 20(1). https://doi.org/10.1145/3365571

Markus, H., & Nurius, P. (1986). Possible selves. American Psychologist, 41(9), 954–969.

McCluney, C. L., Durkee, M. I., Smith, R. E., Robotham, K. J., & Lee, S. S.-L. (2021). To be, or not to be…Black: The effects of racial codeswitching on perceived professionalism in the workplace. Journal of Experimental Social Psychology, 97, 104199. https://doi.org/10.1016/j.jesp.2021.104199

McGee, E. O., & Martin, D. B. (2011). “You Would Not Believe What I Have to Go Through to Prove My Intellectual Value!” Stereotype Management Among Academically Successful Black Mathematics and Engineering Students. American Educational Research Journal, 48(6), 1347–1389. https://doi.org/10.3102/0002831211423972

Meyer, J. H. F., Shanahan, M. P., & Laugksch, R. C. (2005). Students’ conceptions of research: A qualitative and quantitative analysis. Scandinavian Journal of Educational Research, 49(3), 225–244. https://doi.org/10.1080/00313830500109535

Moll, L. C., Amanti, C., Neff, D., & Gonzalez, N. (1992). Funds of knowledge for teaching: Using a qualitative approach to connect homes and classrooms. Theory into Practice, 31(2), 132–141.

National Science Foundation. (2021). Women, Minorities, and Persons with Disabilities in Science and Engineering: 2021 (Special Report NSF 21-321.). National Center for Science and Engineering Statistics. https://ncses.nsf.gov/wmpd.

Potvin, G., & Hazari. (2013). The development and measurement of identity across the physical sciences. Physics Education Research Conference 2013, Portland, OR. https://www.compadre.org/Repository/document/ServeFile.cfm?ID=13182&DocID=3729

Pratt, M. G., Rockmann, K. W., & Kaufmann, J. B. (2006). Constructing professional identity: The role of work and identity learning cycles in the customization of identity among medical residents. Academy of Management Journal, 49, 235–262. https://doi.org/10.5465/AMJ.2006.20786060

Remich, R., Naffziger-Hirsch, M. E., Gazley, J. L., & McGee, R. (2016). Scientific growth and identity development during a postbaccalaureate program: Results from a multisite qualitative study. CBE—Life Sciences Education, 15(3), ar25.

Rincón, V., & Hollis, L. (2020). Cultural code-switching and Chicana/O post-secondary student persistence: A hermeneutic phenomenological analysis. Journal of Latinos and Education, 19(3), 232–245. https://doi.org/10.1080/15348431.2018.1499516

Robnett, R. D., Chemers, M. M., & Zurbriggen, E. L. (2015). Longitudinal associations among undergraduates’ research experience, self-efficacy, and identity. Journal of Research in Science Teaching, 52(6), 847–867.

Robnett, R. D., Nelson, P. A., Zurbriggen, E. L., Crosby, F. J., & Chemers, M. M. (2018). Research mentoring and scientist identity: Insights from undergraduates and their mentors. International Journal of STEM Education, 5(1), 41. https://doi.org/10.1186/s40594-018-0139-y

Rodriguez, S. L., Perez, R. J., & Schulz, J. M. (2022). How STEM lab settings influence graduate school socialization and climate for students of color. Journal of Diversity in Higher Education, 15(1), 58.

Saldaña, J. (2016). The coding manual for qualitative researchers (3rd ed.). Sage Publications.

Schinske, J. N., Perkins, H., Snyder, A., & Wyer, M. (2016). Scientist spotlight homework assignments shift students’ stereotypes of scientists and enhance science identity in a diverse introductory science class. CBE—Life Sciences Education, 15(3), ar47. https://doi.org/10.1187/cbe.16-01-0002

Shiffman, S., Stone, A. A., & Hufford, M. R. (2008). Ecological Momentary Assessment. Annual Review of Clinical Psychology, 4(1), 1–32. https://doi.org/10.1146/annurev.clinpsy.3.022806.091415

Stanton, J. D., Means, D. R., Babatola, O., Osondu, C., Oni, O., & Mekonnen, B. (2022). Drawing on internal strengths and creating spaces for growth: How black science majors navigate the racial climate at a predominantly white institution to succeed. CBE—Life Sciences Education, 21(1), ar3. https://doi.org/10.1187/cbe.21-02-0049

Stone, A. A., & Shiffman, S. (1994). Ecological momentary assessment (EMA) in behavorial medicine. Annals of Behavioral Medicine, 16, 199–202. https://doi.org/10.1093/abm/16.3.199

Suddaby, R. (2006). From the editors: What grounded theory is not. Academy of Management Journal, 49(4), 633–642. https://doi.org/10.5465/amj.2006.22083020

Thiry, H., Laursen, S. L., & Hunter, A.-B. (2011). What experiences help students become scientists? A comparative study of research and other sources of personal and professional gains for STEM undergraduates. The Journal of Higher Education, 82(4), 357–388.

Thiry, H., Weston, T. J., Laursen, S. L., & Hunter, A.-B. (2012). The benefits of multi-year research experiences: Differences in novice and experienced students’ reported gains from undergraduate research. CBE—Life Sciences Education, 11(3), 260–272. https://doi.org/10.1187/cbe.11-11-0098

Thompson, J. J., & Jensen-Ryan, D. (2018). Becoming a “science person”: Faculty recognition and the development of cultural capital in the context of undergraduate biology research. CBE—Life Sciences Education, 17(4), ar62. https://doi.org/10.1187/cbe.17-11-0229

Tracy, S. J. (2010). Qualitative quality: Eight “big-tent” criteria for excellent qualitative research. Qualitative Inquiry, 16(10), 837–851. https://doi.org/10.1177/1077800410383121

Tufford, L., & Newman, P. (2012). Bracketing in qualitative research. Qualitative Social Work, 11(1), 80–96. https://doi.org/10.1177/1473325010368316

Tuma, T. T., Adams, J. D., Hultquist, B. C., & Dolan, E. L. (2021). The dark side of development: A systems characterization of the negative mentoring experiences of doctoral students. CBE—Life Sciences Education, 20(2), ar16.

van Veelen, R., Derks, B., & Endedijk, M. D. (2019). Double trouble: How being outnumbered and negatively stereotyped threatens career outcomes of women in STEM. Frontiers in Psychology, 10. https://www.frontiersin.org/articles/10.3389/fpsyg.2019.00150

Vasquez-Salgado, Y., Camacho, T. C., López, I., Chavira, G., Saetermoe, C. L., & Khachikian, C. (2023). “I definitely feel like a scientist”: Exploring science identity trajectories among Latinx students in a critical race theory-informed undergraduate research experience. Infant and Child Development, n/a(n/a), e2371. https://doi.org/10.1002/icd.2371

Verderame, M. F., Freedman, V. H., Kozlowski, L. M., & McCormack, W. T. (2018). Point of view: Competency-based assessment for the training of PhD students and early-career scientists. ELife, 7, e34801. https://doi.org/10.7554/eLife.34801

Vough, H. C., Caza, B. B., & Maitlis, S. (2020). Making sense of myself: Exploring the relationship between identity and sensemaking. In A. D. Brown (Ed.), The Oxford Handbook of Identities in Organizations (online edn, pp. 244–260). Oxford University Press. https://doi.org/10.1093/oxfordhb/9780198827115.001.0001

Walton, G. M. (2014). The new science of wise psychological interventions. Current Directions in Psychological Science, 23(1), 73–82. https://doi.org/10.1177/0963721413512856

Walton, G. M., & Wilson, T. D. (2018). Wise interventions: Psychological remedies for social and personal problems. Psychological Review, 125(5), 617–655.

Watts, R. J., Diemer, M. A., & Voight, A. M. (2011). Critical consciousness: Current status and future directions. New Directions for Child and Adolescent Development, 2011(134), 43–57. https://doi.org/10.1002/cd.310

Weiner, B. (1985). An attributional theory of achievement motivation and emotion. Psychological Review, 92(4), 548–573.

Yosso, T. J. (2005). Whose culture has capital? A critical race theory discussion of community cultural wealth. Race Ethnicity and Education, 8(1), 69–91.

Zuckerman, A. L., & Lo, S. M. (2022). Examining the variations in undergraduate students’ conceptions of successful researchers: A phenomenographic study. CBE—Life Sciences Education, 21(3), ar55. https://doi.org/10.1187/cbe.21-10-0295

