## Additional Information for "Beyond performance, competence, and recognition: Forging a science researcher identity in the context of research training"

***Corresponding Author:**

Mariel A. Pfeifer

Additional Information File Inventory



| File Description | Page |
| --- | --- |
| Additional File 1. Screening questions for Phase 2 interviews | 2-3 |
| Additional File 2. Interview questions for Phase 2 | 4-5 |
| Additional File 3. Description of themes and categories with example data | 6-11 |
| Additional File 4. Extended positionality statement of research team | 12-13 |
| References | 14 |

**Additional File 1**. Screening and demographic questions for Phase 2 interviews.

**Screening questions**

1. What university or institution are you affiliated with?

2. Have you completed at least two years of an undergraduate degree in the natural sciences (physics, chemistry, geology, biology, and associated subfields)?

3. In one to two sentences, summarize what type of research you do.

4. Choose from the options below
    a. I am a current undergraduate student with at least 2 months of research experience as an intern in a faculty member's research group.
    b. I am a current postbaccalaureate researcher with at least 2 months of research experience as an intern in a faculty member's research group.
    c. I am a current PhD student.
    d. None of these (If this option selected, participant asked to summarize their current academic level and how long they have been in their current research experience in text).

5. If participant selected 4C…Select the best description of your current status in your PhD program.
    a. I am a first-year student who has been in my current research lab for 6 months to one year.
    b. I am a second or third-year student who completed my PhD candidacy requirements within the last 6 months.
    c. I am a fourth-year or higher graduate student who is one to two-years away from completing my degree program.
    d. None of these. (If this option selected, participant asked to describe current status in PhD program in text).

**Demographic questions**

6. Are you the first person in your family to attend college?
    a. Yes
    b. No
    c. Don’t know
    d. Prefer not to answer

7. With which racial or ethnic group(s) do you identify? Please choose all that apply.
    a. Black or African American
    b. American Indian or Alaskan Native
    c. Latina/Latino/Hispanic
    d. Native Hawaiian or Other Pacific
    e. White
    f. None of these (If this option selected, participant asked to describe what racial or ethnic groups they identify in text).
    g. Prefer not to answer

8. Which gender do you most identify with?
    a. Woman
    b. Man
    c. Nonbinary/third gender
    d. Transgender
    e. Prefer not to say

**Additional File 2**. Interview protocol for Phase 2. Questions related to a separate study are redacted.

Introductory comments and verbal consent
Thanks for scheduling an interview with me, and for responding to our survey about your research experiences. You agreed to participate in this study when you responded to the survey, but I want to check if you have any questions before we proceed with the interview portion. Today's interview will be audio and video recorded. We will transcribe your comments and replace any identifying information with pseudonyms. We will only use the transcript in our study, so no one else will hear your voice or know that you participated in the study. *Do you have any questions for me at this point?*

Proceed with questions. Start recording.

1. Tell me about yourself. How do you describe yourself to someone new?

2. What does a typical day of research "work" look like for you right now?
    a. Follow-up: How is this work done?
    b. Follow-up: Tell me about your interactions with your research supervisors.
    c. Follow-up: Tell me about your interactions with your peers.

3. How would you describe a science researcher to a friend who is not involved in science?

4. Do you think the work you do is the work of a science researcher? Why or why not?

5. Is there work you do now that you do not consider to be the work of a science researcher? If so, why?

6. Do you think you can be yourself when you do science research? Why or why not?

7. Do you see yourself, today, as a science researcher? Why or why not?
    a. Follow-up: Is there anything (previous experiences, other people, outside activities, etc.,) that helps you feel like a science researcher?
    b. Follow-up: How do these experiences, people, or activities help you see yourself as a science researcher?

8. Can you think of a time / situation when you didn't see yourself as a science researcher? Tell me about that…

9. How, if at all, has your view of science research changed because of your research experience?
    a. If there is a change, follow-up: Does this change in knowledge influence how you see yourself as a science researcher?

10. Has your view of who science researchers are changed since you began doing research? If so, how?
    a. If there is a change, follow-up: Does this change in knowledge influence how you see yourself as a science researcher?

**11.** Do you think you have changed as a person because of your current research experience? Why or why not?
   a. If yes, follow-up: What are these changes? What caused these changes?

**12.** How, if at all, does the culture of your research group, or lab, influence how you see yourself as a science researcher?
   a. (Define culture as values, beliefs, norms, if participant asks).
   b. Follow-up: Who in your lab is considered a science researcher? Why?
   c. Follow-up: How, if at all, does the talk in your lab influence how you see yourself?

**13.** What are your career goals? Are you interested in doing science research as part of this career? Why or why not?

**14.** Is there anything else I should know about your research experience and how it relates to you seeing yourself as a science researcher?

Generic follow ups used throughout interview.
You mention _______, tell me more about that.
You mention _______, can you give me an example of what that looks like for you?
You mention _______, what was that like for you?

**Additional file 3**. Summary of emergent findings with example quotes from Phase 1 and Phase 2 data.

| *Theme* | *Category* | *Description* | *Phase 1 Example* | *Phase 2 Example* |
|---|---|---|---|---|
| Recognizing oneself as a science researcher<br><br>(Carlone & Johnson, 2007) | Science student | ECRs recognizing themselves as a "student" and not yet a researcher. Includes thinking that the purpose of their work is to learn how to become a researcher. | I feel like a student doing research rather than a scientist. | I would say when I started working in a lab and I was an undergrad just taking classes and most of what I was doing in the lab was washing bottles. I don't think then I considered myself a scientist. I was a science student.<br>-Iris<br>[*overlaps with nature of work*] |
| | Science researcher | ECRs recognizing themselves as a "researcher" with the purpose of their work being testing hypotheses or addressing research questions. | Now, I think of myself as a scientist because I showed myself that I can solve a research question. | I do see myself as a science researcher because well, my daily task is to [make] progress on a question that needs to be answered. -Seth |
| | Career researcher | ECRs describing the work of a career researcher as securing funding, setting big-picture research goals, and managing team of researchers. | *Not observed in dataset* | And if I can be called a science researcher, and [my PI] could be called a professional [researcher], our lives look a lot different. They're managing the lab and keeping it afloat, doing a lot of administrative stuff, writing grants, reviews, keeping up with the literature, and looking at the things that we're doing and trying to synthesize them and know where everyone's at. [They have] the big picture in mind. Whereas, for me, if I'm a [science] researcher, my day-to-day is just I'm down in the weeds. I'm not worrying about trying to write grants…My job is pretty much like, you have questions that you need to answer. -Andrew |

| *Theme* | *Category* | *Description* | *Phase 1 Example* | *Phase 2 Example* |
| --- | --- | --- | --- | --- |
| Recognizing oneself as a science researcher, continued | Epistemic involvement<br><br>(Burgin et al., 2012) | Intellectual responsibility for research. | Negative example<br>In a lot of ways, I feel less like a scientist because much of the work I do is just tasks for graduate students. I don't often know why I am doing it or get to see the data afterward, so it doesn't feel like I am actually being a scientist. I would feel more like a scientist if I knew more about the projects, my contributions, and the outcomes. | Negative example<br>I do participate in research, but in terms of like the actual assay development and the thought processes behind this stuff, I'm not as involved as I was in my previous research [experiences] [in] undergrad. [There] I had full autonomy on my project. But at my current job, I basically just, process samples and just get stuff done. I don't have to think too deeply about what I do outside of how to be consistent and how to make sure things are in place so that people can do their jobs… I'm not involved in the decision making for the projects.<br>-Justin |
| | Operational autonomy<br><br>(Bailyn, 1985) | Discretion to select the means to achieve a particular research task or goal. | This research experience has made me feel more like a scientist because never before have I been given this much trust to do an experiment correctly without supervision. | [As an undergraduate researcher] I was kind of helping someone else. Someone else was guiding me and providing help. [Now] I have to decide what I'm doing on a day-to-day basis. I get to decide what time I come into work and when I leave. -Lisa |
| | Strategic autonomy<br><br>(Bailyn, 1985) | Independence in determining research goals and projects to pursue. | *Not observed in dataset* | I will feel more like a science researcher when I get to the point of fully creating my own project and answering questions from scratch, because up to this point, I still need some guidance. I think that is really common in [starting graduate students], especially. I was never going to walk in and be like, "I want to do this." Especially with my first project, it's building off of others. I have this dataset and a lot of direction.. where I get to the point where I can direct my own research, regardless of what it is, I think that's when I'll feel like a researcher.-Claire |

| *Theme* | *Category* | *Description* | *Phase 1 Example* | *Phase 2 Example* |
|---|---|---|---|---|
| Making sense of influential factors | Perceived cohesion<br><br>(Bollen & Hoyle, 1990) | Cognitive and/or affective appraisal of relationship to the research community, that can range from a local research community to the larger academic research community. | I don't feel like I belong since I don't relate to anybody else who works in the lab. | I just felt [like] you probably are not meant to be here [doing research in this place], maybe this is not for you… -Ezra, *continues below in critical reflection* |
| | Attributions<br><br>(Weiner, 1985) | Causal thinking used to interpret or understand one's own science identity and how it is affected, especially apparent when an individual experiences a negative social interaction. | This is somewhat hard to completely gage... One of my friends that has different lab partners had a much more collaborative experience than me, which is what I identify as something that would probably have allowed me to feel like a scientist. But I didn't have that experience, so I felt like [research] was mostly busy work that I did on my own. | From the very start of undergrad, I felt like I could see myself as a scientist or a science researcher… That has pretty much stayed constant. Like the bad experiences I had [in a previous lab] mostly just made me mad. I did not think that the problem was me.<br>-Talia, sharing why she perceived her science identity was protected when she felt limited perceived cohesion within a previous research group |
| | Critical reflection<br><br>(Watts et al., 2011) | Acknowledgement that systemic problems within science research such as racism, sexism, classism, etc. contribute to diminished perceived cohesion and/or recognizing oneself as a science researcher. | *Not observed in dataset* | (*continued from above in perceived cohesion*)…But once I worked on that and changed the way I thought about it. Now, it's like, no, you got to make sure that everyone has an opportunity in this type of environment. It's [Research is] not just meant for a [single] group of people. -Ezra |

| *Theme* | *Category* | *Description* | *Phase 1 Examples* | *Phase 2 Example* |
|---|---|---|---|---|
| Individual factors | Research and researcher conceptions | Thoughts, beliefs and attitudes about researchers and research, including conceptions of qualifications to be a researcher. | I feel like I'm not a real scientist until I get my PhD. | I feel like we kind of all grow up with this idea that science is done by one genius person who makes a cool discovery and then goes on and wins a Noble Prize, doing a bunch of work by themself, but that's absolutely false. Science is collaborative. Everybody is making small steps with the help of other people, with the help of the work of the people who came before… So you really feel like, yeah, you don't need to be a really super smart genius to do scientific research and to make a positive impact on science. You can just be you…-Cindy |
| | Skill perceptions<br><br>(Carlone & Johnson, 2007) | Assessment of own research abilities (insufficient or sufficient), including how ECRs remind themselves that they do possess sufficient research skills to recognize themselves as a researcher. | <u>Insufficient</u><br>[I feel] less like a scientist because everything I did in the lab failed. None of my mutagenic PCRs were successful and neither was my PAGE.<br><br><u>Sufficient</u><br>This research experience has made me feel more like a scientist because I feel that I now have the skills and resources necessary to perform well in the field of research. I have learned a lot about scientific writing, laboratory procedure, and how to collaborate with my peers among other things. | But I'll just kind of look down on myself or just [think] maybe you're not good at this [research]. What I've done is I've edited my CV over the last few years…And what I'll do is, I'll go look back at previous versions of my CV. And look to my CV now. And it's like, you have done a lot…Look at all this on my CV. Like I have done all this…I am a science researcher.-Max |

| *Theme* | *Category* | *Description* | *Phase 1 Examples* | *Phase 2 Example* |
|---|---|---|---|---|
| Individual factors, continued | Career intentions<br><br>(Hazari et al., 2010; Markus & Nurius, 1986) | Plans for future job and how participant sees research as part of that job (or not), including assessment of why they think they would be happy or successful in a certain type of career, which relates to possible selves. Possible selves are future versions of oneself related to being a science researcher, including both positive and negative possible selves. | I am still interested in research questions and topics, especially in biology and kinesiology. However, this class has confirmed my thoughts that I would not enjoy the day-to-day life of only being a researcher. I plan to only pursue research as a part of graduate school then work in sports medicine. These feelings about general research are from the structure and environment I [had] in this [experience]. | And working with [my PI], I mean they're very productive. They get up very early to start doing work. And at least from my perspective of it, they work until they've gone to bed, basically. And to me, that's not something that I, anymore, want. -Ross |
| Contextual factors | Nature of work<br><br>(Pratt et al., 2006) | The topic of research or focus of daily work and how the research tasks are accomplished affecting science identity. | My research experience was less scientific (for example, we did not work in a lab looking at cell cultures), so it is hard to say that it made me feel more like a scientist. | [In one of my research labs] I did field work with them. Like I was like out in the field, like on my knees, digging up dirt, getting bit by a thousand mosquitoes. Interacting with the things that I am studying. So [my work for that project] has really enabled me to develop that relationship with my work and with the people in the lab. So that experience, that definitely informs how I feel as a scientist. Like I feel like it really genuinely gives me more authority on things. Cause I have like one more step of understanding how this [system] works because I'm out there looking at it.-Mitchell |

| *Theme* | *Category* | *Description* | *Phase 1 Examples* | *Phase 2 Example* |
|---|---|---|---|---|
| | Social interactions<br><br>(Carlone & Johnson, 2007) | Dealings with other individuals in a science context that influence how an individual recognizes themselves as a science researcher. These interactions can be negative, absent, or positive. Negative and absent interactions result in an individual questioning or doubting their science identity and positive interactions support science identity. | I would say more like a scientist, but this is because of the way I was treated by my supervisor more than the work I did in the lab. My supervisor was very encouraging and often thanked us for our hard work and assured us that we were doing something important. | *See data in main text in social interactions section of results and discussion.* |
| | Group norms<br><br>(Pratt et al., 2006) | How the implicit, expected patterns behavior or standard ways of operating within a context an ECR is embedded within influences their science identity, career intentions and/or possible selves. Contexts include research groups, programs, departments, institutions, and disciplines within natural sciences. | Being exposed to what it's like to work in a research lab outside of assigned class work is really interesting. It gives a better picture of what it would be like as a career and more of what a typical day looks like beyond the broad picture of scientists that the media portrays. | Now I'm in this lab where it's like super flashy… my lab mates, they're on NPR [National Public Radio]… [The notoriety of the lab] makes [research] feel even more serious and like more high stakes... your lab group can shape how you see yourself as a researcher.- Skylar |

**Additional File 4**. Extended positionality statement of research team.

While analyzing data, we considered our own privileges and how they may affect our interpretations. We acknowledge that as a research team we identified as white, which may have limited our interpretations. We endeavored to address the limitation by engaging in a form of member checking and by seeking additional feedback on emergent results from individuals outside our own research team. The positionality of each author is described below.

The first author (M.A.P.) served as the lead researcher for this project. She is a postdoctoral researcher associated with a science and technology center with research experience in two research fields (discipline-based science education research and cell biology). Throughout her training, she often experienced doubt in seeing herself as a researcher, in part because she perceived the knowledge, skills, and abilities acquired in one research field were not necessarily viewed by meaningful individuals in the other field as assets. She further identifies as a white, cis-gendered woman from a first-generation, lower middle-class, rural background and felt resonance with ECRs who explained that they did not see themselves as a science researcher because they did not know anyone in their community growing up who was a researcher.

The second author (C.J.Z.) is a graduate student pursuing a doctoral degree in biochemistry and molecular biology with a research focus on science education. He holds bachelors' degrees in biology and earth sciences and has research experience in environmental and planetary sciences as well as discipline-based science education. He identifies as a white, cis-gendered man from a continuing generation, middle-class, suburban background. Because he experienced identity tensions when switching from one field of research to another field of research, he found he could relate to ECRs who were facing a similar field-to-field transition. He felt he could find common ground with ECRs who described their concerns of not being an integral part of their research teams due to his own previous research experiences.

At the time of data analysis, the third author (J.M.I.) recently completed a bachelor's degree in biology and was intending to pursue a master's degree in environmental management. He identifies as a white, cis-gendered man from a continuing-generation, upper-class, suburban background. He participated in undergraduate research for a total of three terms before engaging in data analysis for this project. Often, he felt a connection with ECRs who discussed how issues related to the nature of work within a specific discipline shaped their identity as science researchers, as well as their future career aspirations.

At the time of data analysis, the fourth author (O.A.E.) recently completed a bachelor's degree in biology and was preparing to begin her first year of medical school. Before beginning data analysis for this project, she had seven terms of research experience. She identifies as a white, cis-gendered, continuing-generation woman from an upper-middle class, suburban background. She provided insights into the experiences of students utilizing research to explore their scientific professional identities and interpretations of comments from an undergraduate's student perspective.

The last author (E.L.D.) is a faculty member in a biochemistry and molecular biology department with a research focus on science education. She holds a bachelor's degrees in biology and a doctoral degree in neuroscience. She has ~20 years of experience conducting discipline-based education research. She identifies as a white, cis-gendered woman from a continuing generation, suburban family. Growing up, her family was middle class and she now considers herself upper-middle-class. She came to recognize the challenges of becoming a science researcher and the privilege of her own continuing-generation researcher status when she was a graduate student. These experiences prompted her to study research training environments as contexts for students' career development and decision making, particularly in the life sciences. She felt like she could bring her knowledge of research and theory related to research training, career development, and equity in STEM education to facilitate understanding of the results and their implications and implications.

### References


Bailyn, L. (1985). Autonomy in the industrial R&D lab. *Human Resource Management*, *24*(2), 129–146. https://doi.org/10.1002/hrm.3930240204

Bollen, K. A., & Hoyle, R. H. (1990). Perceived cohesion: A conceptual and empirical examination. *Social Forces*, *69*(2), 479–504. https://doi.org/10.2307/2579670

Burgin, S. R., Sadler, T. D., & Koroly, M. J. (2012). High school student participation in scientific research apprenticeships: Variation in and relationships among student experiences and outcomes. *Research in Science Education*, *42*(3), 439–467. https://doi.org/10.1007/s11165-010-9205-2

Carlone, H. B., & Johnson, A. (2007). Understanding the science experiences of successful women of color: Science identity as an analytic lens. *Journal of Research in Science Teaching*, *44*(8), 1187–1218. https://doi.org/10.1002/tea.20237

Hazari, Z., Sonnert, G., Sadler, P. M., & Shanahan, M.-C. (2010). Connecting high school physics experiences, outcome expectations, physics identity, and physics career choice: A gender study. *Journal of Research in Science Teaching*, *47*(8), 978–1003. https://doi.org/10.1002/tea.20363

Markus, H., & Nurius, P. (1986). Possible selves. *American Psychologist*, *41*(9), 954–969.

Pratt, M. G., Rockmann, K. W., & Kaufmann, J. B. (2006). Constructing professional identity: The role of work and identity learning cycles in the customization of identity among medical residents. *Academy of Management Journal*, *49*, 235–262. https://doi.org/10.5465/AMJ.2006.20786060

Watts, R. J., Diemer, M. A., & Voight, A. M. (2011). Critical consciousness: Current status and future directions. *New Directions for Child and Adolescent Development*, *2011*(134), 43–57. https://doi.org/10.1002/cd.310

Weiner, B. (1985). An attributional theory of achievement motivation and emotion. *Psychological Review*, *92*(4), 548–573.
